# Timing the Evolution of Phosphorus-Cycling Enzymes Through Geological Time

**DOI:** 10.1101/2023.08.21.554175

**Authors:** Joanne S. Boden, Juntao Zhong, Rika E. Anderson, Eva E. Stüeken

## Abstract

Phosphorus plays a crucial role in controlling biological productivity, but geological estimates of phosphate concentrations in the Precambrian ocean, during life’s origin and early evolution, vary over several orders of magnitude^1-10^. While reduced phosphorus species may have served as alternative substrates to phosphate^11^, their bioavailability on the early Earth remains unknown. Here, we reconstruct the phylogenomic record of life on Earth and find that phosphate transporting genes (*pnas*) evolved in the mid-Archean (ca. 3.2 Ga) and are consistent with phosphate concentrations above modern levels (>3 μM). The first gene optimized for low phosphate levels (*pstS*; <1 μM) appeared around 2.9 Ga. Most enzymatic pathways for metabolising reduced phosphorus emerged and expanded across the tree of life in the Neoarchean to Paleoproterozoic (ca. 2.6 to 1.8 Ga). This includes phosphonate-catabolising CP-lyases, phosphite-oxidising pathways and hypophosphite-oxidising pathways. CP-lyases are particularly abundant in dissolved phosphate concentrations below 0.1 μM. Our results thus indicate declining phosphate levels through the Archean, possibly linked to increasing deposition of phosphate-scavenging iron oxides, which may have limited productivity. However, reduced phosphorus species did not become widely used until after the Paleoproterozoic Great Oxidation Event (2.3 Ga), possibly linked to an expansion of the biosphere at that time.

## INTRODUCTION

Phosphorus is one of the fundamental building blocks of life. It is an essential constituent of DNA, cell membranes and ATP, but is quickly depleted by plants and microbes in marine and terrestrial settings worldwide. As a result, restricted phosphorus availability limits primary productivity in large areas of the continents and oceans ^12,13^.

It is conceivable that Precambrian microbial communities were similarly limited by phosphorus, and if so, that may have constrained the growth and evolution of the biosphere. However, geochemical estimates of the Precambrian marine phosphorus reservoir vary by several orders of magnitude, from less than 0.12 μM to 4,100 μM in the Archean and between 0.03 and 2.5 μM in the Proterozoic oceans ^1-10^. It has also been proposed that local microscale environments could have achieved up to 100 mM orthophosphate to drive prebiotic chemical pathways ^14^. These disparities arise from differences in sampling strategies and assumptions in computational models. Therefore, the extent to which phosphorus was available to microbes in the Archean and Proterozoic remains a topic of debate, especially with regards to surface waters where fewer direct constraints exist.

Modern microbial communities cope with limited phosphorus availability using a rich variety of enzymes to source phosphorus from the environment, including a range of dissolved organic phosphorus (DOP) compounds and inorganic phosphorus species, collectively referred to as total dissolved phosphorus (TDP) (Figure 1). These include orthophosphate and phosphate esters (where P is in a fully oxidized state; P(V)) as well as more reduced compounds in the form of phosphite (P(III)), phosphonates (likely P(III)) and hypophosphite (P(I))^15-23^. Phosphite and hypophosphite constitute up to 26% of DOP in some habitats ^21^ and geochemical estimates suggest that phosphite in particular, was prominent in the Paleoarchean ocean due to elevated input from meteorites and Earth’s more reducing redox state ^24^. Furthermore, phosphite is a source of phosphorus for multiple microbial species ^19^, and in some cases it can serve as an electron donor for chemolithotrophs ^22,25,26^. In contrast, phosphonates make up approximately a quarter of all high-molecular weight DOP in the global oceans ^27^. They contain carbon chains and characteristically strong carbon-phosphorus (C-P) bonds which are not present in other reduced phosphorus molecules. They can be assimilated as a nutrient resource by a diverse range of bacterial and archaeal taxa across the tree of life and in several different habitats ^15,16,18-20^. Importantly, genomic studies have found that phosphonate-catabolising pathways catalysed by CP-lyases, phosphonatases, and some dioxygenases are more common in phosphate-depleted waters ^17,28^ and soils ^29^, suggesting that they may represent a mechanism for coping with phosphate scarcity. This observation implies that the phylogenetic history of these enzymes can be used as a tool for reconstructing phosphate availability in the past.

**Figure 1:**
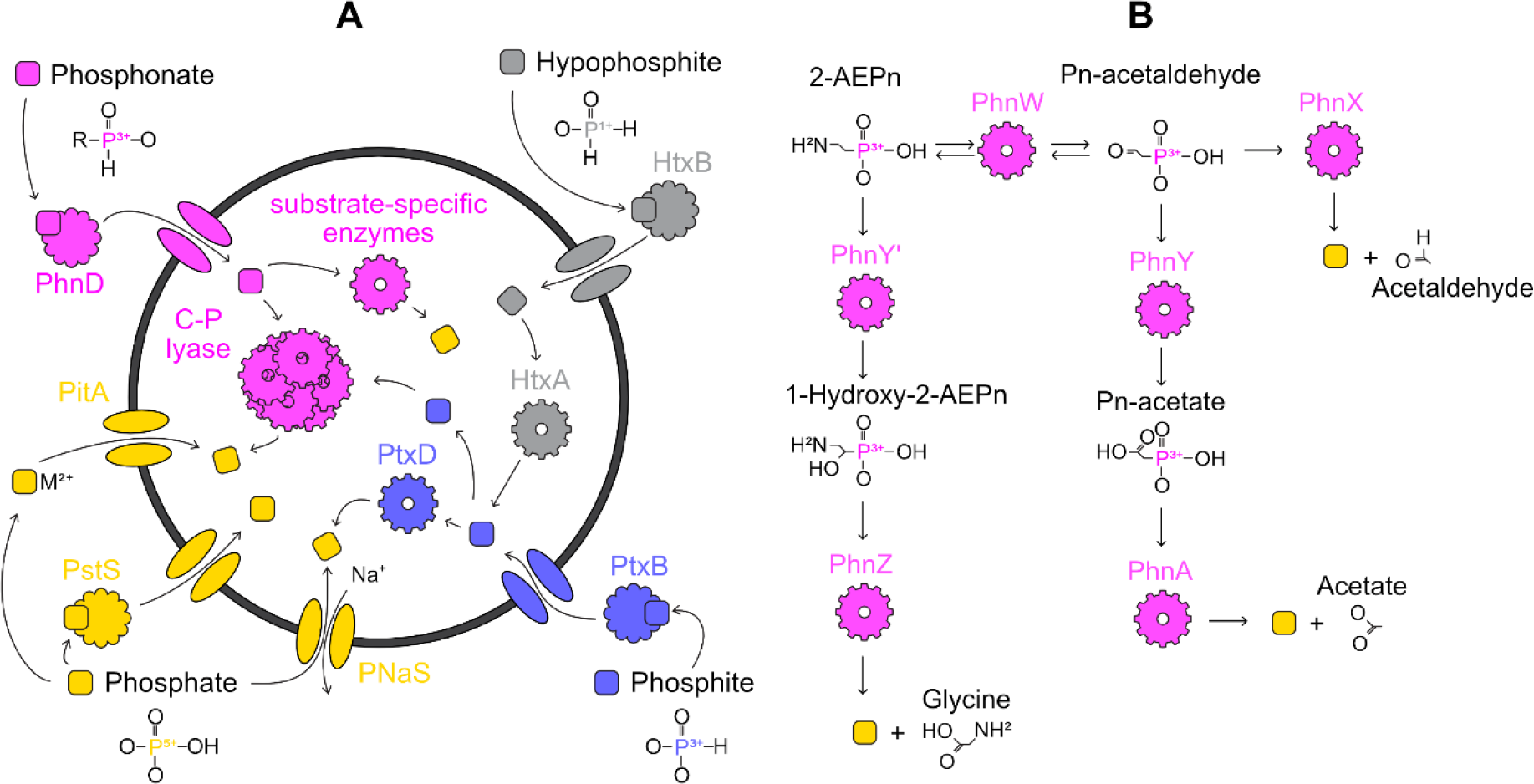
Enzymatic pathways for importing and catabolising reduced phosphorus molecules and phosphate. An overview of the metabolisms researched in this project (A) and a detailed summary of substrate-specific phosphonate degradation pathways (B). Coloured squares indicate hypophosphite (grey), phosphite (blue), phosphonate (pink) and phosphate (yellow) molecules, whereas other shapes of the same colour indicate the enzymes responsible for catalysing their import and catabolism.

Here, we perform phylogenetic analyses to investigate which phosphorus compounds were available for microbial communities over geologic time. More specifically, we reconstruct the evolutionary history of microbial genes involved in utilising phosphate, phosphonates, phosphites, and hypophosphites. To do this, we reconcile a time-calibrated tree of life with phylogenetic trees of phosphorus-utilizing genes to estimate when these genes were gained, lost, duplicated, and vertically inherited (see Methods). We then infer past phosphate concentrations by comparing presence/absence and abundance patterns for each gene with phosphate concentrations in the modern ocean. Our results allow us to shed new light on the evolution of the biogeochemical phosphorus cycle and provide the first constraints on the availability of reduced phosphorus substrates.

## RESULTS

### Tree of Life

To determine how ancestral microbes used the various phosphorus species considered in this study, we began by reconstructing a tree of life from 865 genomes, chosen to represent the full diversity of life and genomes harbouring *ptxD*. Our results are broadly consistent with previous studies (e.g. ^30-34^), because Eukaryotes are sisters of Asgaard Archaea (ultrafast bootstrap support (UBS) = 21), and DPANN are the deepest branches of the archaeal subtree (UBS = 100). Similarly, Planctomycetes, Chlamydia and Verrucomicrobia are closely related to each other (UBS = 91) ^31,34-36^.

To estimate when key groups of organisms evolved, we implemented one uncorrelated (UGAM) and two autocorrelated (LN and CIR) Bayesian molecular clock models on alignments of 16 ribosomal proteins that have been used previously to reconstruct the tree of life ^30^. The resulting time-calibrated phylogenies made with CIR, LN and UGAM clock models suggest that the last universal common ancestor radiated into Bacteria and Archaea at ca. 4.35 Ga (confidence intervals range from 4.40-4.22 Ga), 4.38 Ga (confidence intervals range from 4.40-4.34 Ga) or 4.37 Ga (confidence intervals range from 4.40-4.29 Ga), respectively. These dates fall shortly after the first evidence for liquid water ^37,38^, which was applied as an upper constraint for the origin of life.

All three molecular clock models also predict that crown bacteria emerged at least 200 million years before crown Archaea (Table 1), consistent with previous studies ^31^.

**Table 1:** Molecular clock estimates timing the emergence of crown bacteria and archaea under three different clock models. Confidence intervals are provided in brackets.

| Clock Model | Bacteria / Ga | Archaea / Ga |
| --- | --- | --- |
| CIR | 3.47 (3.61 to 3.31) | 3.26 (3.45 to 3.07) |
| LN | 3.80 (3.91 to 3.67) | 3.26 (3.45 to 3.08) |
| UGAM | 3.90 (4.28 to 3.57) | 3.32 (3.51 to 3.13) |

### Distribution of Phosphorus-Cycling Genes in the Tree of Life

We surveyed 865 genomes for 13 genes involved in the utilisation and production of reduced phosphorus molecules and phosphate. The results reveal a broad distribution of phosphate sodium symporters (namely *pnas*) and active phosphate transport mechanisms (indicated by *pstS*) throughout 257 and 174 strains of our dataset respectively (Figure 2). These encompass all major bacterial and archaeal phyla except DPANN and Patescibacteria (formerly CPR, Figure 2). In contrast, genes for low-affinity phosphate import of orthophosphate ions bound to divalent metal ions, such as Mg^2+^, Ca^2+^, Co^2+^, Mn^2+^ and Zn^2+^ via *pitA* and *pitH* are absent or rare respectively. Only one of the 865 genomes harbours a *pitH* homolog (Figure 2).

**Figure 2:**
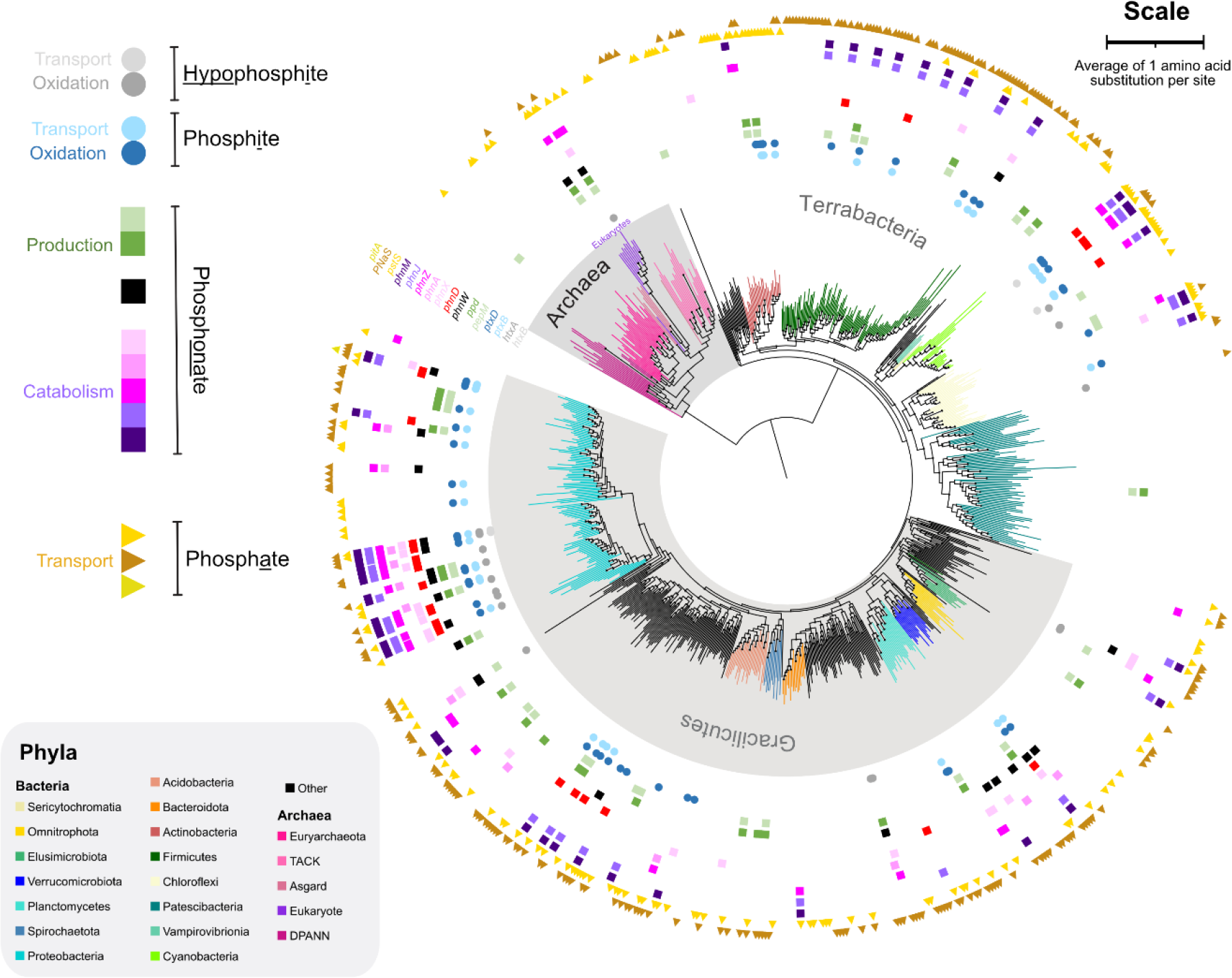
Distribution of phosphorus-cycling genes in the tree of life. Coloured shapes outside branch tips represent homologs of enzymes involved in hypophosphite, phosphite (circles), phosphonate (squares) and phosphate (triangles) utilisation and production. Branches are coloured according to their phylum. The evolutionary tree was reconstructed from 16 ribosomal proteins using maximum likelihood methodology. The scale bar represents an average of one amino acid substitution per site.

Reduced phosphorous metabolisms are present in a variety of bacteria, but most archaea cannot metabolise phosphonates, phosphite or hypophosphite (Figure 2). Of these, bacteria from the Proteobacterial superphylum host a particularly broad variety of phosphonate-cycling genes, as has been documented in previous studies ^17,18^. We also find that phosphonate production is rarer than its catabolism, (consistent with previous work ^15^), because only 30 genomes harbour *pepM* and *ppd* to produce phosphonates from phosphate esters, whereas 94 genomes have at least one of *phnX, phnA, phnZ, phnJ* or *phnM* to catabolise phosphonates. Of all the phosphonate catabolism genes, *phnJ* and *phnM* encoding the broad-specificity CP-lyases are most common because they co-occur in 44 genomes compared to the substrate-specific phosphonate catabolism genes, *phnX, phnW, phnA* and *phnZ* which are each present in just 28, 25, 12 and 40 genomes respectively (Extended Data – Table 1). The distribution of these genomes in the tree of life (Figure 2) reveals a broader taxonomic distribution of phosphonate-producing compared to phosphonate-catabolising organisms, which is consistent with previous work ^17^.

Hypophosphite is more reduced than phosphonates and phosphite. As a result, only 4 genomes have *htxB* encoding the binding protein required for active hypophosphite import from the environment. Two of these genomes also have *htxA* for oxidising hypophosphite to phosphite, but a further 15 genomes have *htxA* without the binding protein for hypophosphite import (Figure 2). It is possible that these *htxA* genes encode enzymes that oxidise formate instead, as biochemical analyses have found comparable rates of activity on these small organic molecules ^39^.

### Timing of Gene Events

To estimate when these phosphorus-cycling genes emerged and radiated across the tree of life, our molecular clocks were reconciled with Bayesian evolutionary trees of each phosphorus-cycling gene to ascertain the timing of speciation, duplication, loss, and horizontal gene transfer events. The most ancient event for a given gene represents the first phylogenetic evidence of that gene’s presence, whereas a clustering of events in a particular time interval represents widespread use of the metabolism across diverse lineages. Any lineages which experienced gene events but subsequently became extinct, do not leave signatures in the genomic record, so in these cases, the origin of each gene could be earlier than we report. Our estimates represent lower bounds.

Estimates from different clock models vary, with UGAM predicting generally younger events than LN and CIR, but all three suggest that microbial communities began importing phosphate before they evolved methods of assimilating reduced phosphorus molecules in the form of phosphonate, phosphite and hypophosphite (Supplementary Information Figure 1A-N). For example, the earliest evidence of the phosphate-sodium symporter *pnas* appears at the Paleoarchean-to-Mesoarchean boundary, and the earliest evidence for a key component of active high-affinity phosphate transporters, *pstS*, emerges in the Neoarchean (Figure 3). For the rest of this paper, we focus on estimates from the CIR clock model, because it has been shown to outperform other clock models ^40,41^. A full report of results from all three clock models in the Supplementary Information.

**Figure 3:**
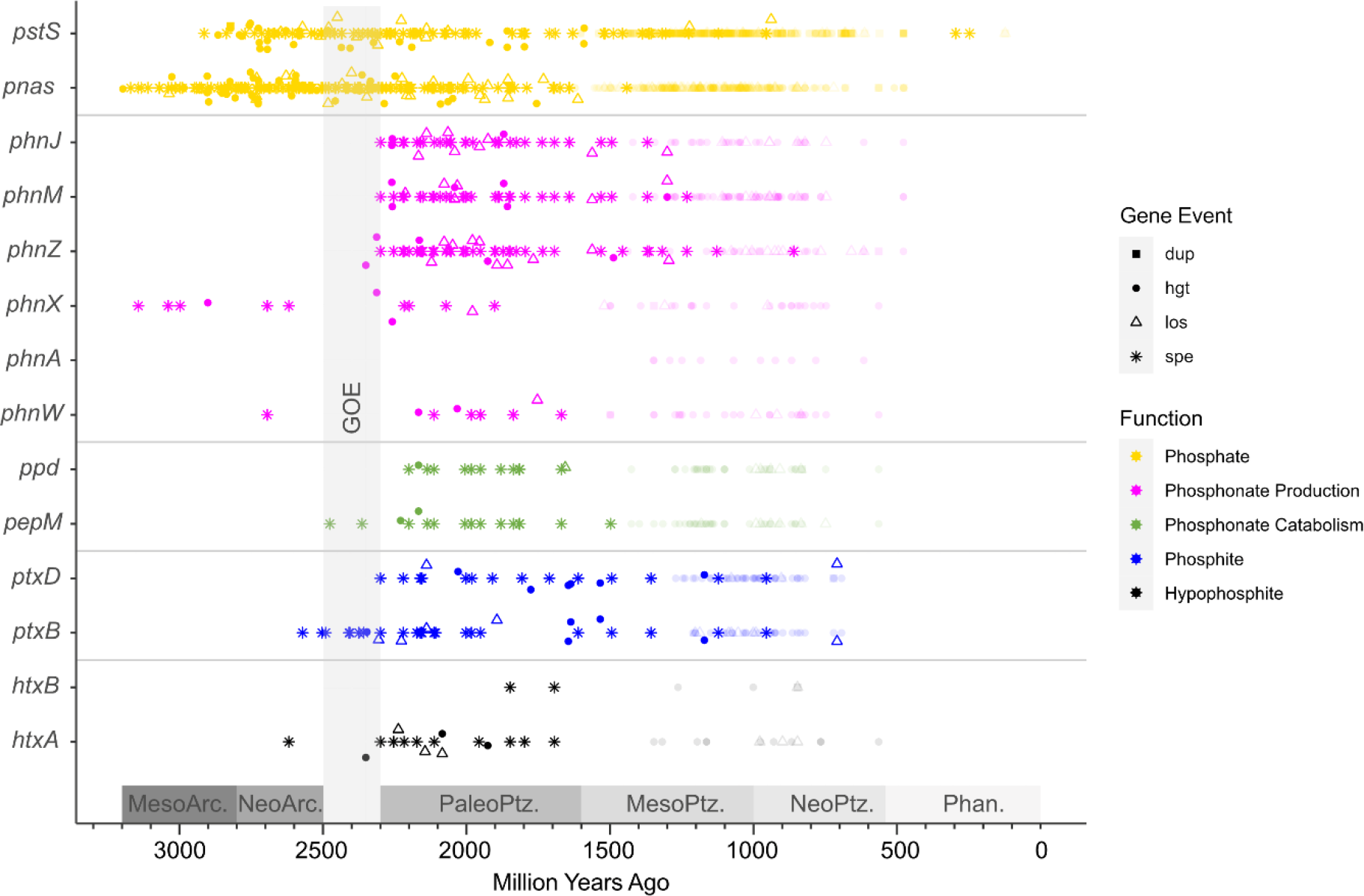
Estimated origin of phosphorus-cycling genes in the tree of life. Coloured shapes represent speciations (asterisks), horizontal gene transfers (filled circles), duplications (filled squares) and losses (empty triangles) of genes underlying phosphate utilisation (yellow), phosphonate usage (pink), phosphonate production (green), phosphite usage (blue) and hypophosphite usage (black). Each transfer, duplication and loss is estimated to have occurred on either an internal (bright colours) or terminal (faded colours) branch of our time-calibrated-tree-of-life, made using the Cox-Ingersoll-Ross clock model. Our methods do not predict whether transfers, duplications and losses happened at the start or end of the branch, so we summarise them by plotting the midpoints. For a full representation of branch lengths and uncertainties, see supplementary information figure 1A-N.

Microorganisms began catabolising phosphonates with the evolution of *phnX* in the Mesoarchean, but the ability remained confined to a small subset of lineages in comparison to phosphate import mechanisms (Figure 3). The gene *phnX* encodes phosphonatase, which acts on phosphonoacetaldehyde to produce phosphate and small organic molecules for downstream metabolisms within the cell (Figure 1). Later, in the Neoarchean, the number of potential phosphonate molecules that could be used as a source of phosphorus increased with the emergence of *phnW*, encoding 2-aminoethylphosphonate-pyruvate transaminase, which can act in concert with *phnX* to release phosphate from 2-aminoethylphosphonate (2-AEP) (Figure 3, Figure 1). The genomic repertoire for catabolizing phosphonates increased in the Paleoproterozoic around 2.3 Ga with the emergence of the genes *phnJ* and *phnM*, encoding broad-specificity-CP-lyases, and *phnZ*, encoding an Fe(II)-dependent phosphohydrolase. These genes would have enabled micro-organisms to liberate phosphate and glycine from a post-cursor of 2-AEP and to catabolise a wide variety of phosphonates (Figure 3).

Biological methods of producing these phosphonates began to emerge in the late Neoarchean, with the evolution of *pepM* for transforming the relatively oxidised phosphate ester phosphoenolpyruvate (P in +5 oxidation state) to the more reduced phosphonate phosphonopyruvate (P in +3 oxidation state) and vice versa. This reaction is reversible, so another enzyme is required to force the reaction in the direction of phosphonate production; *ppd* (otherwise known as *aepY*) encodes phosphonopyruvate decarboxylase, which has been found to be one of the most common methods of doing this^16^. Here, we find phylogenomic evidence to suggest that *ppd* emerged after *pepM* in the early Paleoproterozoic (Figure 3).

The earliest evidence for microbial utilisation of phosphite and hypophosphite appeared in the Neoarchean with the advent of *ptxB* for importing phosphite and *htxA* for catalysing the oxidation of hypophosphite and formate (Figure 3)^39^. It has previously been hypothesised that dissimilatory phosphite oxidation (DPO) emerged in the most recent common ancestor of Firmicutes and Deltaproteobacteria ^22^, which emerged **∼** 3.2 Ga ^42^ or **∼** 2.9 Ga ^43^ depending on which molecular clocks are used to infer their age. Here, we predict that this ancestor arose 3.45 Ga (confidence intervals span 3.30-3.59 Ga), but our gene-tree-species-tree-reconciliations do not find evidence of *ptxD* in this ancestor. Instead, the earliest event indicative of the presence of *ptxD* is a speciation which occurred within the Firmicutes 2.30 Ga (confidence intervals span 2.56 to 2.22 Ga, Figure 3). This places an upper limit on the origin of DPO in the early Paleoproterozoic. However, *ptxB* encoding part of an active phosphite importer evolved earlier, so phosphite could have been imported for use in other biological processes in the late Neoarchean.

## DISCUSSION

Our results allow us to place new constraints on phosphate concentrations in the Precambrian ocean. Many reduced-phosphorus cycling genes are only present in microbial communities when more easily accessible forms of phosphorus, such as orthophosphate, are not available ^17,19,28,29^. We performed regression analyses of data from the *Tara* Oceans Project ^44,45^ and found significant correlations between the abundance of four genes in our dataset and phosphate concentrations (Figure 4). Gene abundances of *pstS, phnM* and *phnJ* reduce with increasing phosphate concentrations (coefficients are −0.79, −0.29 and −0.32 respectively), whereas the abundance of *nptA* from the *pnas* symporter family decreases (coefficient = 4.78 x 10^-5^, Figure 4). For the remaining nine genes, the relevant data are not available. These trends imply that genes encoding CP-lyases and high affinity phosphate transporters are present in low phosphate environments and relatively rare in regions with phosphate concentrations approaching 3.29 μM (Figure 4). Similar negative correlations between marine phosphate concentrations and phosphonate-cycling genes, including CP-lyases, phosphonatase (encoded by *phnX*) and Fe(II)-dependent phosphonohydrolases (encoded by *phnZ*) have been documented previously in global surface oceans ^17,28^ and terrestrial soils ^29^. In contrast, *nptA* which is a member of the low affinity PNaS symporter family, is more abundant in regions with high phosphate concentrations (> 1 μM) (Figure 4).

**Figure 4:**
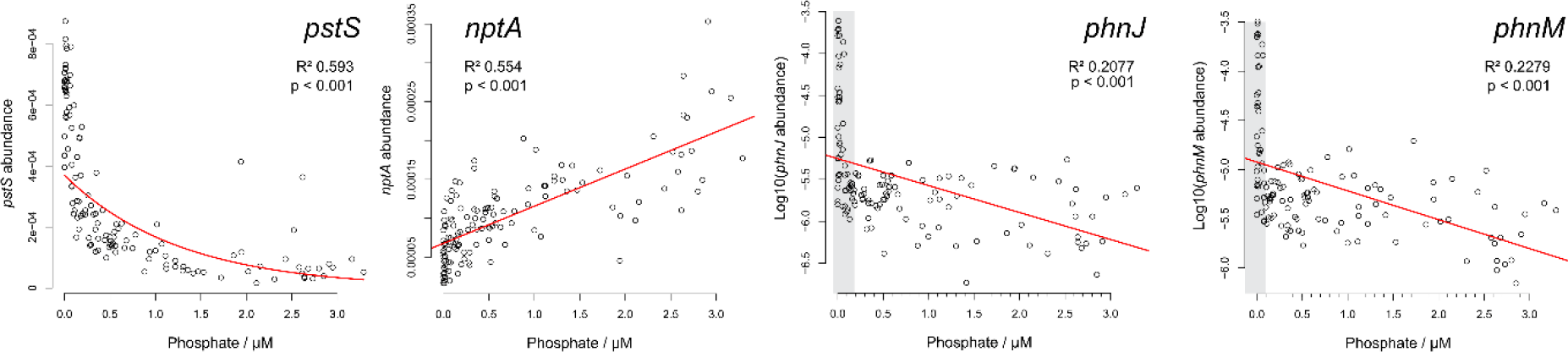
Regression analyses of phosphorus-cycling genes in regions of the global ocean with varying phosphate concentrations. The abundance of reads containing *pstS, nptA, phnJ* and *phnM* are plotted on the y axis against micromolar phosphate concentrations on the x-axis, for all 139 samples from the Ocean Microbial Reference Catalog v2 (OM-RGC.v2) of the *Tara* Oceans Project ^44,45^. Regression lines are shown in red. Grey areas indicate low phosphate concentrations where the abundances of CP-lyase genes are particularly high.

These correlations, combined with our estimates on the origin of phosphate transporters and broad-specificity CP-lyases can be used to provide estimates on phosphate availability in the Archean and Paleoproterozoic. Geochemical studies have proposed dissolved phosphate concentrations ranging from less than 0.004 ^2-5^ up to 4,100 μM ^1^ in the Archean ocean, which is up to **∼** 1,000 times higher than the highest phosphate concentration in the *Tara* dataset (0 to 3.29 μM). Given the positive linear correlation of the sodium-phosphate symporter, *nptA*, with phosphate availability (Figure 4), it is likely that higher than modern phosphate concentrations were present on Earth during the early Mesoarchean when the first evidence of enzymes from each family appears (Figure 3), perhaps consistent with the estimates of Ingalls, et al. ^7^ and Rasmussen, et al. ^6^.

Toward the end of the Mesoarchean, more areas of the planet must have experienced low phosphate concentrations to drive the emergence of more metabolically-expensive methods of obtaining phosphorus using *phnX* and the high-affinity phosphate transporter *pstS* (Figure 3). Extrapolations of our exponential regressions suggest that at 1,000 μM phosphate, *pstS* would be extremely scarce (Figure 4). At 25 μM phosphate, *pstS* abundance would also be very low, which would probably not lead to a strong selective pressure driving the evolution of the gene (Figure 4). As a result, we consider it more likely that late Mesoarchean organisms (Figure 3) would have evolved *pstS* in areas with low phosphate concentrations of 1 μM or less, in line with geochemical estimates by several studies ^2,3,5,9,10,46^. By extension, areas with low phosphate must have existed throughout the Neoarchean, Proterozoic and Precambrian to drive the speciation, duplication, and gene transfer events of *pstS* that are predicted in our molecular clocks (Figure 3).

The relationships of *phnJ* and *phnM* with phosphate provide further insight into Paleoproterozoic environments, where they first emerged (Figure 3). These genes enable micro-organisms to use phosphonates as a source of phosphorus and small organic molecules. Therefore, their abundances exhibit a stark drop-off above 0.1 μM phosphate (Figure 4). Above this threshold the highest abundances of *phnM* and *phnJ* drop by an order of magnitude and median values drop by a factor of 4 to 5. As a result, environments with low phosphate concentrations (below 0.1 μM) are likely to have driven the evolution of broad-specificity phosphonate catabolism in the Paleoproterozoic. This is within the range predicted by some geological records of Paleoproterozoic deep ocean habitats (e.g. ^4,10^ and ^3^), but lower than others (e.g. ^5^).

In order to understand whether our estimates of phosphate availability are relevant for terrestrial or marine environments of the Paleoproterozoic and Archean, we need to know where the ancestral organisms which harboured *pnas, pstS, phnJ* and *phnM* were living. Some insight can be gained from our inferred gene events. Our reconciliations predict that the Archean speciations of *pstS* and *pnas* were numerous and occurred in a range of bacterial phyla, including Armatimonadota, Bacteroidota, Patescibacteria, Actinobacteria, Chloroflexi, Omnitrophota, Acidobacteriota, Thermoplasmatota, Cyanobacteria and Firmicutes (Extended Data Figure 1). In contrast, Paleoproterozoic speciations of *phnJ* and *phnM* were rarer and occurred in Verrucomicrobia, Firmicutes and Proteobacteria (Extended Data Figure 2). Members of these phyla today occupy many different habitats, including the terrestrial subsurface ^47^ and marine water column ^48^. As a result, it is possible that a large range of habitats exhibited similar or lower than modern phosphate concentrations in the Neoarchean and Paleoproterozoic.

## CONCLUSIONS

By tracing the evolutionary trajectories of enzymes for importing and utilising a range of phosphorus species, we have provided insight into the microbial usage of phosphorus compounds throughout the Precambrian. The early Mesoarchean emergence of a phosphate sodium importer (*pnas)* that is most abundant in phosphate-replete environments, followed by phosphonatase and a high affinity phosphate transport binding protein (*pstS)* that are most abundant in phosphate-deplete environments (Figures 3 and 4)^17^, suggests that the phosphate concentrations available to many microbes dropped from the middle to the end of the Archean (Figure 5). This drop would be consistent with increased deposition of banded iron formations (BIF) in the Neoarchean compared to the Mesoarchean ^49^. BIFs are composed of ferric iron oxides, which have a high affinity for phosphate adsorption and may therefore have scavenged phosphate from the water column ^9,10^. In the absence of iron oxidation, all iron would have been in the Fe(II) state, which renders phosphate relatively more soluble, possibly allowing phosphate concentrations of hundreds to thousands of μM ^1^.

**Figure 5:**
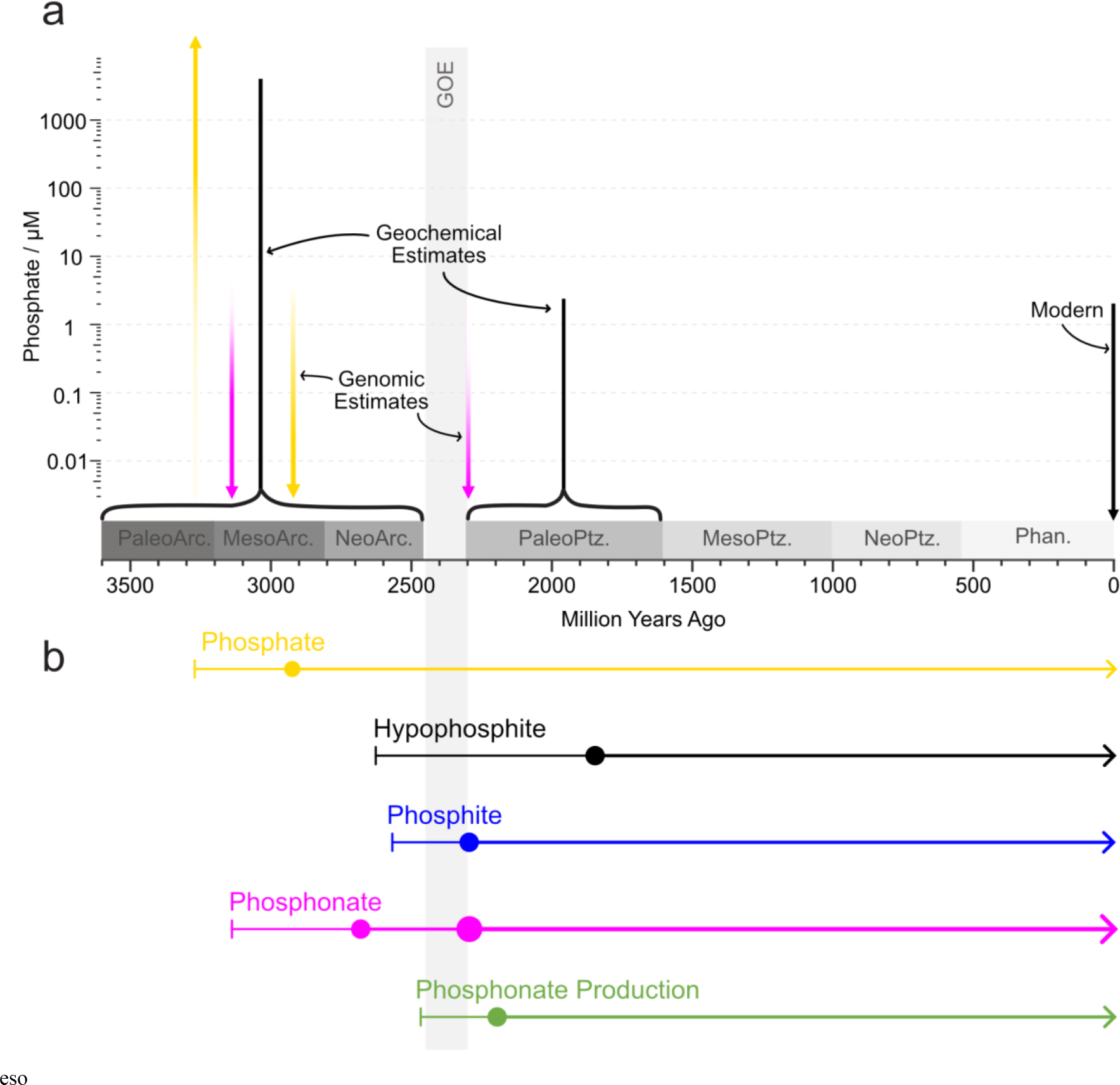
Estimates on the availability of orthophosphate (a) based on genomic records tracing the origin of phosphorus-cycling enzymes (b). Geochemical estimates of marine phosphate concentrations (vertical black bars) for the Archean and Proterozoic are indicated alongside phosphate concentrations most likely to have driven the evolution of the phosphate transport enzymes PstS and PNaS (vertical yellow arrows) and the phosphonate-metabolising enzymes PhnJ and PhnM (pink bars). Coloured horizontal lines represent the time since enzymes involved in the microbial utilisation of phosphate (yellow), hypophosphite (black), phosphite (blue) and phosphonate (pink) are estimated to have existed. Green horizontal lines represent the time since enzymes which produce phosphonates have existed. Circles represent the evolution of subsequent enzymes involved in the utilisation or production of each molecule. GOE, Great Oxygenation Event.

A drop in phosphate from the early to late Mesoarchean could have limited biological productivity and O_2_ production by Cyanobacteria, which perhaps delayed the rise of pO_2_ in Earth’s atmosphere, i.e. the Great Oxidation Event (GOE), to 2.45 Ga, long after the evolutionary origin of oxygenic photosynthesis. Indeed, our limited analyses of 22 cyanobacterial genomes included in the molecular clocks and reconciliations predict that Cyanobacteria were genomically capable of using phosphate sodium symporters in the early Neoarchean, and only acquired *pstS* for active phosphate import toward the end of the Neoarchean (Extended Data Figure 1).

Whilst multiple microbial species from different phyla were genomically capable of importing phosphate in the Archean, fewer species could use reduced phosphorus compounds (as indicated by the relative scarcity of events in Figure 3). This appears to have changed at the Neoarchean to Paleoproterozoic boundary, when phylogenetic evidence points toward the evolution of genomic mechanisms for assimilating phosphite (via *ptxD*) and a broad variety of phosphonate molecules (via *phnJ* and *phnM*). This post-GOE expansion in the phosphorus pool available to microbes may be a consequence of the growing biosphere ^46,50^. Some biogeochemical models suggest that primary productivity was limited by the availability of electron donors until the evolution of oxygenic photosynthesis, which became widespread at the start of the GOE ^50^. Similarly, models of ancient carbon cycles suggest that marine net primary production increased by two orders of magnitude from the Neoarchean to Paleoproterozoic ^46^. As microbial communities grew, more and more organisms would have competed for the available phosphorus, perhaps increasing the selective pressures driving the evolution of genomic mechanisms for utilising alternative phosphorus species.

The evolution of biological phosphonate production with the emergence of *ppd* in the Paleoproterozoic may also have been a result of more crowded microbial habitats as phosphonates can act as competitive inhibitors of phosphate and carboxylic acid functional groups in metabolic pathways, interrupting the growth of other microorganisms ^15^. They are more resistant to degradation than phosphate esters, owing to their stable C-P bonds, which has led to the hypothesis that marine microbes produce them to protect against viral infection and grazing ^15^.

In conclusion, our phylogenetic analyses provide insight into the bioavailability of phosphorus molecules and the concentrations of phosphate throughout the Archean and Proterozoic. We provide the first estimates on the origin of alternative phosphorus species and use an independent approach to estimate Precambrian phosphate concentrations around microbial communities. Together, these suggest that phosphite and hypophosphite were not widely used by the ancestors of present-day biota for most of the Archean. Phosphonate use dates back to the Mesoarchean, but like phosphite and hypophosphite it appears to have become more widely used after the GOE, possibly linked to biological diversification and increased competition. Lastly, our results advance ongoing debates about the Precambrian phosphate reservoir and help reconcile previously conflicting geochemical estimates. Phosphate-replete conditions inferred from solubility experiments ^1^ are consistent with genomic data for the early Mesoarchean, whereas phosphate-depleted conditions proposed by others ^2,3,5,9,10,46^ may be linked to increased BIF deposition in the Neoarchean, as supported by the emergence of high-affinity phosphate transporters at that time. Phosphate limitation may therefore have been a significant controlling factor in the timing of the GOE.

## MATERIALS AND METHODS

### Species Tree Reconstruction

Our evolutionary tree of life was constructed from information present in the ribosomal proteins of 865 genomes, which were chosen to represent the full diversity of bacteria and archaea using methods described in Mateos, et al. ^51^. In order to achieve this, we included one representative from each order of the tree of life using information from the Genome Taxonomy Database (GTDB) release 95 ^52,53^. Where possible, a representative with the gene *ptxD* was included to better capture the diversity of phosphite-oxidising organisms. From these genomes a species tree was reconstructed using 16 ribosomal proteins (namely L18, L3, L5, S8, L4, S3_C, L6, L2, L15e, S19, S17, L22, S10, L24, L16 and L14) that have been used previously to infer the tree of life ^30,54,55^. Their homologs were obtained from each genome with HMMER3 v3.3.2 ^56^(95 % of the genomes contained homologs of eight or more of these genes), then aligned and trimmed with Muscle v5.1 ^57^ and TrimAl v1.4.rev15 ^58^, all implemented in GToTree v1.6.34 ^59^.

The resulting alignments were used to reconstruct a species tree using maximum likelihood methodology implemented in IQTREE v2.0.3 ^60^. Partitioned analyses ^61^ were applied to allow each protein to evolve under appropriate substitution models. To find these substitution models, ModelFinder was implemented with default options (no mixture models) and ‘MERGE’ ^62^. The best was found to be a single partition containing all proteins, so the LG substitution rate matrix was applied to all 16 ribosomal protein alignments, with 10 categories of the FreeRate model to estimate substitution rates and their variations at different sites. Further variations in substitution rates through time (as described in Lopez et al ^63^) were modelled by applying the −Q option to account for heterotachy.

Information from previous studies which thoroughly investigated macroevolutionary relationships between large clades of bacteria and archaea was incorporated by applying a topological constraint on the relationships between Gracilicutes, Terrabacteria, Patescibacteria/CPR and Euryarchaeota in our tree (Extended Data Figure 3). For all analyses, branch supports were estimated with 1,000 replicates of ultrafast bootstrap approximations ^64^ and the SH-aLRT test ^65^. Three independent replicates were completed, but only one consensus tree with the best (a.k.a. closest to zero) log-likelihood value was retained for further analyses.

### Divergence Time Estimation

Our species tree was anchored to a geological timeline by implementing a relaxed Bayesian molecular clock in Phylobayes v.4.1^66^. Topology was fixed to match our species tree, and ages estimated based on predicted mutation rates for the 16 ribosomal proteins described above (2,290 aligned amino acid positions). Substitutions were modelled using a uniform exchangeability matrix and an empirical profile mixture model with 20 profiles (specified using -catfix C20) ^67^. Divergence times were estimated under one of three models to reflect differing views on the inheritance of substitution rates between mother and daughter lineages: These were the autocorrelated lognormal (LN) and Cox-Ingersoll-Ross (CIR) models where substitution rates are inherited from parent lineages ^41,68^, and the uncorrelated gamma multipliers (UGAM) model where substitution rates are chosen independent of ancestry ^69^.

The first diversification in our clock, representing the first radiation of the last universal common ancestor of life on Earth (LUCA) into bacteria and archaea, was set by applying a uniform distribution spanning 4.4-3.5 Ga to the root node. This is based on the earliest evidence for liquid water on the planet’s surface ^37,38^ and some of the earliest generally accepted microfossils ^70^.

We also implemented eight calibration points based on microfossils of cyanobacteria and algae and geochemical evidence for methanogenesis and oxygenic photosynthesis (Extended Data Table 2). Many of these have been applied previously (e.g. ^51,71-77^, but this is the first time they have been combined in a single molecular clock (Extended Data Figure 4).

Two independent chains were run for 43,650 to 156,348 cycles, and were considered converged when the relative difference of all parameters was > 0.3 and the effective sizes of all parameters were > 50. This was assessed using tracecomp implemented in phylobayes v4.1 ^66^, discarding 10,914 to 49,182 cycles as burn-in. A suitable burn-in was chosen based on traces viewed in Tracer v.1.7.0 ^78^.

### Identification of Phosphorus-Cycling Genes

We searched in our molecular clock genome taxa, for homologs of genes involved in key pathways for the degradation, transport and production of phosphonates, phosphite and hypophosphite using HMMER3 ^56^. To do this, we used HMM profiles downloaded from the NCBI’s prokaryotic genome annotation pipeline (Extended Data Table 3) and an e value threshold of 0.1. Where these were unavailable (e.g. for proteins associated with phosphite and hypophosphite transport and oxidation), searches were conducted using the same e value threshold and custom HMM profiles built from experimentally-characterised proteins (Extended Data Table 4) that were aligned in MAFFT v. 7.4 ^79^ with L-INS-I.

The resulting hits were retrieved and aligned with MAFFT v.7.4 ^79^ using E-large-INS-1. Next, they were trimmed to remove positions composed primarily of to ensure (>85%) with trimAl v1.2rev59 ^58^. This cut-off was chosen because less aggressive trimming has been found to better recover well-established relationships ^80^. Maximum likelihood trees were then generated with IQTREE v.2.0.3 ^60^ under substitution models chosen by ModelFinder ^62^ with 1,000 ultrafast bootstraps ^64^.

To the resulting trees, we annotated bitscores from the HMM searches using the ‘Branch score’ and ‘Propagate attribute’ modules in TreeViewer v2.0.1 ^81^. In addition to assigning bitscore values to leaves of the tree, we assigned bitscore values to internal nodes based on the average score of their descendants. Any leaves representing homologs which passed the bitscore thresholds of their respective TIGRFAMs were also highlighted, so that contours could be drawn delimiting clades with the same degree of similarity as the query sequences. Leaves belonging to a monophyletic clade containing the query sequences and any others which passed the respective bitscore thresholds were retained for further analyses, whilst the rest were discarded (Supplementary Information, Figure 2A-N). Similar filtering methods have been employed to identify light-harvesting genes ^73^ and arsenic resistance genes ^82^ in prokaryotes.

### Phylogenetic Analyses of Phosphorus-Cycling Genes

After filtering, the remaining amino acid sequences of each protein family were aligned with MAFFT v. 7.4 ^79^ and trimmed to remove positions comprising >70% gaps with trimAl ^58^. Bayesian phylogenetic trees were reconstructed using MrBayes v. 3.2.7a ^83^ with a mixed amino acid model prior, invariant sites and gamma distributed site rates. Chains were considered converged when the average standard deviation of split frequencies was < 0.01, the potential scale reduction factor between 1.00 and 1.02, and the ESS scores of all parameters were > 200 after the first 25% of iterations were discarded as burn-in. These were calculated with MrBayes and Tracer v1.7 ^78^. Bayesian phylogenetic trees of the two phosphate-transporting enzymes, PstS and PNaS, did not converge within three months, so maximum likelihood methodology was implemented instead, using IQ-TREE v2.0.3 ^60^ with 1,000 ultrafast bootstraps ^64^ and a substitution model chosen by ModelFinder ^62^. To ensure all complex mixture models were included, the following flags were used: -mrate E,I,G,I+G,R and -madd C10,C20,C30,C40,C50,C60,EX2,EX3,EHO,UL2,UL3,EX_EHO,LG4M,LG4X,CF4.

Between 100 and 700 iterations were performed before the ultrafast bootstraps converged. A spot check on phosphonatase revealed that similar results were found regardless of whether MrBayes of IQ-TREE were used to reconstruct the evolutionary trees (Extended Data Figure 5).

### Reconciliation of Gene Trees with the Molecular Clock

To estimate the timing for duplication, transfer, loss, and speciation events for each gene, each gene tree was reconciled with our time-calibrated species tree using ecceTERA v. 1.2.5 ^84^. Default settings were applied (event costs: HGT=3, duplication=2, loss=1, speciation=0; transfer to the dead allowed so HGTs can occur with unsampled lineages) with amalgamate=true to find the most parsimonious reconciled gene tree from a set of gene trees representing uncertainty in their topology ^85^. These gene tree sets were obtained from the .t files of the MrBayes runs using a custom script which discarded the first 25 % as burn-in. From the output of these gene-tree-species-tree reconciliations (specifically the symmetric median reconciliation), we collected the timing of all speciations, transfers, duplications, and losses of each gene by editing Python scripts that were first designed and applied to time the evolution of sulphur-cycling enzymes ^51^.

### Phosphate Concentration

To investigate whether organisms with low-affinity phosphate transporters and CP-lyases are found in environments with high or low phosphate concentrations, we examined the Ocean Microbial Reference Catalog v2 (OM-RGC.v2) from the *Tara* Oceans Project ^44,45^. The OM-RGC.v2 included relative abundances of all clusters of orthologous genes (COG) in 139 *Tara* Oceans metagenomic samples. We associated each gene of interest with a unique COG accession number using the COG search tool (https://www.ncbi.nlm.nih.gov/research/cog/). To quantify how gene abundance varies with phosphate concentration, regression models of the abundance of each COG were fitted to the raw phosphate concentration using the ‘lm’ function of the ‘stats’ package in R. All analyses include 129 degrees of freedom.

## ACKNOWLEDGMENTS

Thank you to our reviewers for their insightful feedback. We also thank Aya Klos and Tony Ni for helpful comments on the manuscript and Aya Klos for helpful conversations regarding R scripting for figures. We also thank Dr. Celine Scornavacca, Dr. Abu Baidya, Dr. Matt Pasek and Dr. Isabelle Bi for helpful discussions. Funding for this work came from a NERC Frontiers grant (NE/V010824/1) awarded to EES. This work was performed by the Virtual Planetary Laboratory Team, a member of the NASA Nexus for Exoplanet System Science, funded via NASA Astrobiology Program Grant No. 80NSSC18K0829.

## AUTHOR CONTRIBUTIONS

E.E.S. R.E.A. and J.S.B conceived the research project; J.S.B and R.E.A. designed the phylogenetic and molecular clock analyses; J.S.B. performed the analyses; J.Z. and J.S.B. found *Tara* data and performed regression analyses; J.S.B, E.E.S. and R.E.A. interpreted the data; all authors wrote and reviewed the manuscript.

## DATA AVAILABILITY

All alignments, protein phylogenies and molecular clocks generated in this study are available in the open science framework repository, https://osf.io/vt5rw/?view_only=b13a53f4d87c44d1a82a18b176523c5b.

## ADDITIONAL INFORMATION

Supplementary Information is available for this paper. Correspondence and requests for materials should be addressed to J.S.B. Reprints and permissions information is available at www.nature.com/reprints.

