## Supplementary Information for "Timing the Evolution of Phosphorus-Cycling Enzymes Through Geological Time"

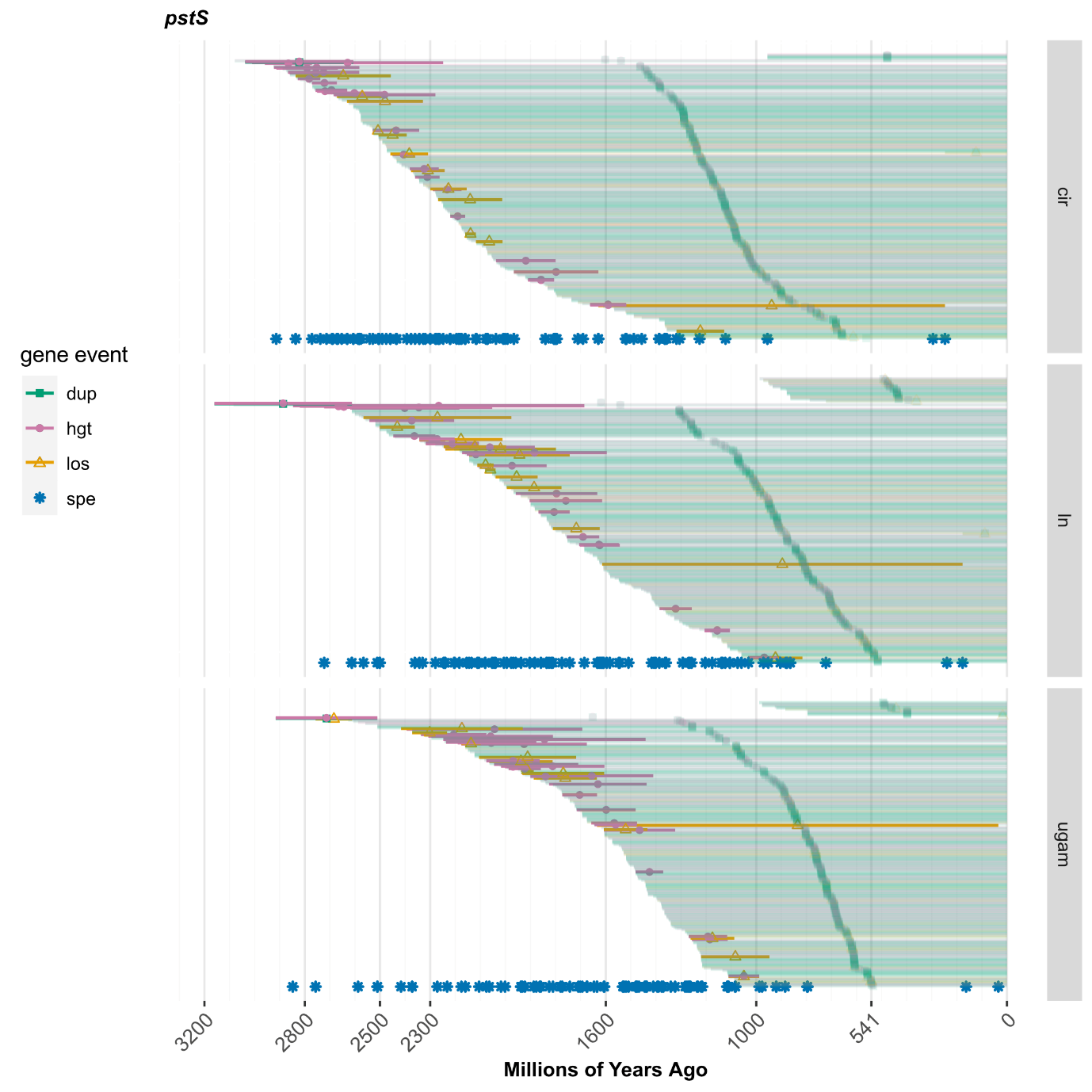

**Figure 1A: Uncertainty in estimating the origin of *pstS*.** Horizontal lines represent the lengths of branches where gene duplications (green), horizontal gene transfers (pink) and losses (orange) are predicted to have occurred. The midpoint of each branch is marked with shapes of the same colour, representing duplications (filled squares), horizontal gene transfers (filled circles) and losses (empty triangles). Darkness indicates whether the event occurred on an internal (dark colour) or terminal (faded) branch of the tree of life. Gene speciations (blue asterisks) are not associated with branch lengths because they occur on internal nodes of the tree. Results found using three different clock models are shown (cir: Cox-Ingersoll-Ross, ln: lognormal, ugam: uncorrelated gamma multipliers).

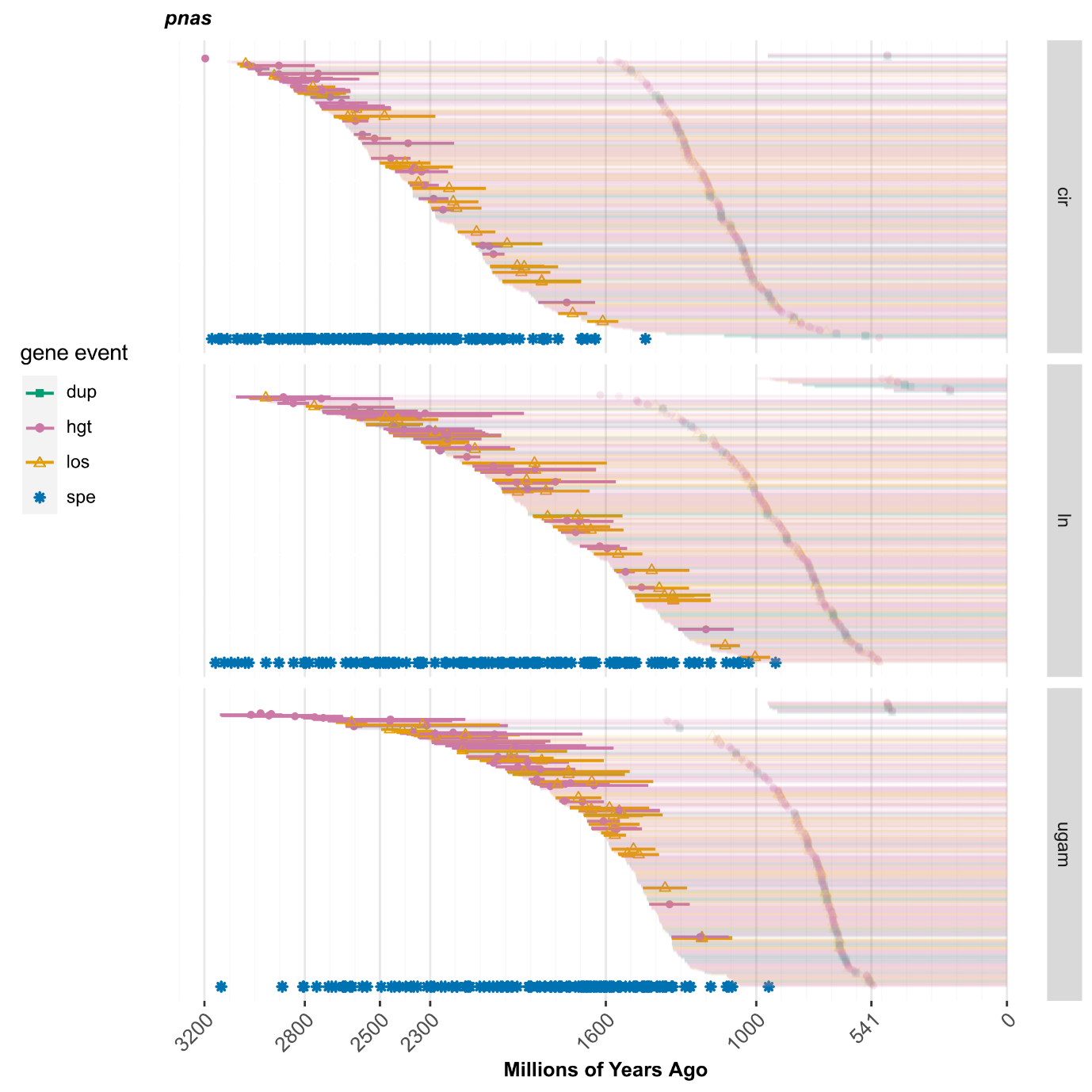

**Figure 1B: Uncertainty in estimating the origin of PNaS.** Horizontal lines represent the lengths of branches where gene duplications (green), horizontal gene transfers (pink) and losses (orange) are predicted to have occurred. The midpoint of each branch is marked with shapes of the same colour, representing duplications (filled squares), horizontal gene transfers (filled circles) and losses (empty triangles). Darkness indicates whether the event occurred on an internal (dark colour) or terminal (faded) branch of the tree of life. Gene speciations (blue asterisks) are not associated with branch lengths because they occur on internal nodes of the tree. Results found using three different clock models are shown (cir: Cox-Ingersoll-Ross, ln: lognormal, ugam: uncorrelated gamma multipliers).

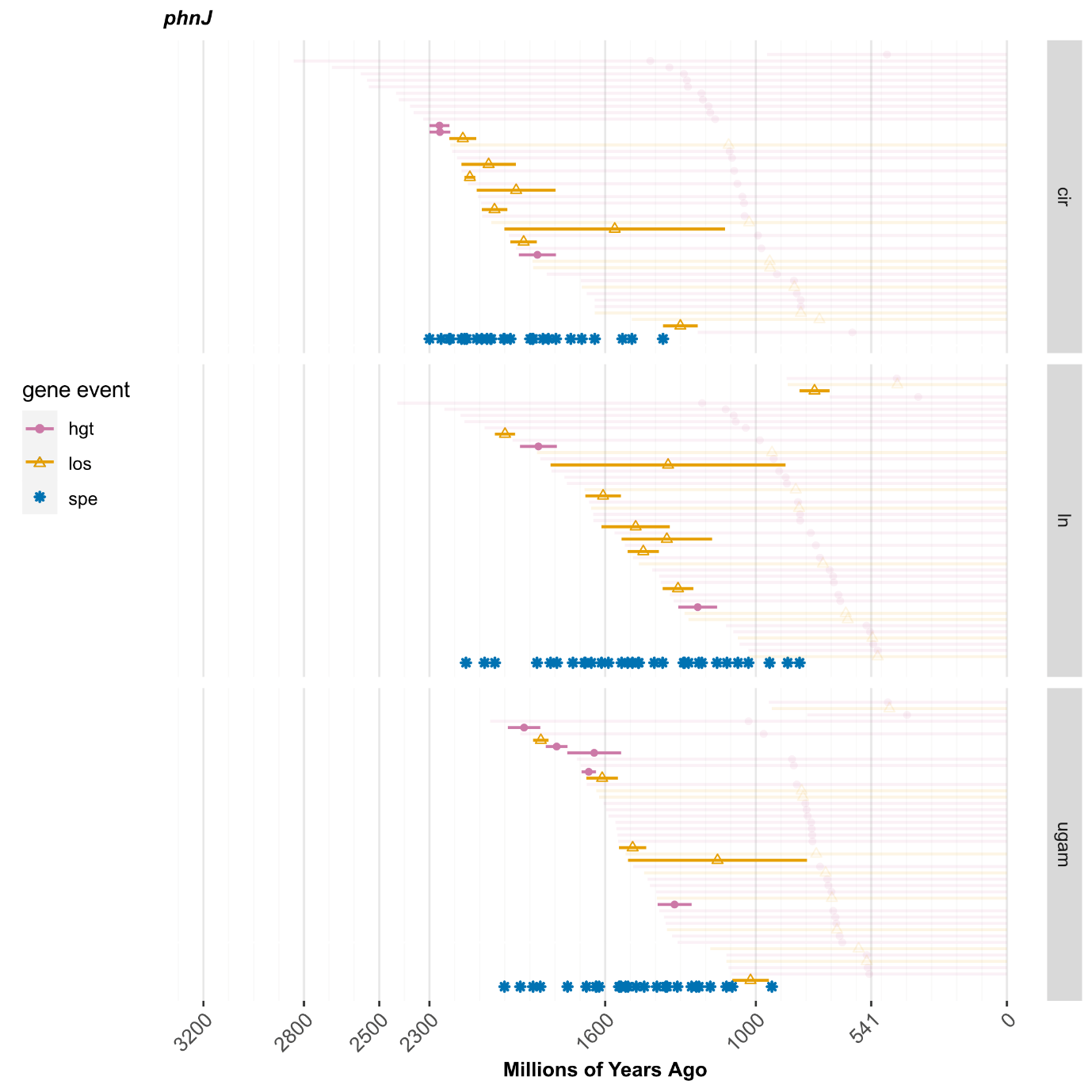

**Figure 1C: Uncertainty in estimating the origin of *phnJ*.** Horizontal lines represent the lengths of branches where gene duplications (green), horizontal gene transfers (pink) and losses (orange) are predicted to have occurred. The midpoint of each branch is marked with shapes of the same colour, representing duplications (filled squares), horizontal gene transfers (filled circles) and losses (empty triangles). Darkness indicates whether the event occurred on an internal (dark colour) or terminal (faded) branch of the tree of life. Gene speciations (blue asterisks) are not associated with branch lengths because they occur on internal nodes of the tree. Results found using three different clock models are shown (cir: Cox-Ingersoll-Ross, ln: lognormal, ugam: uncorrelated gamma multipliers).

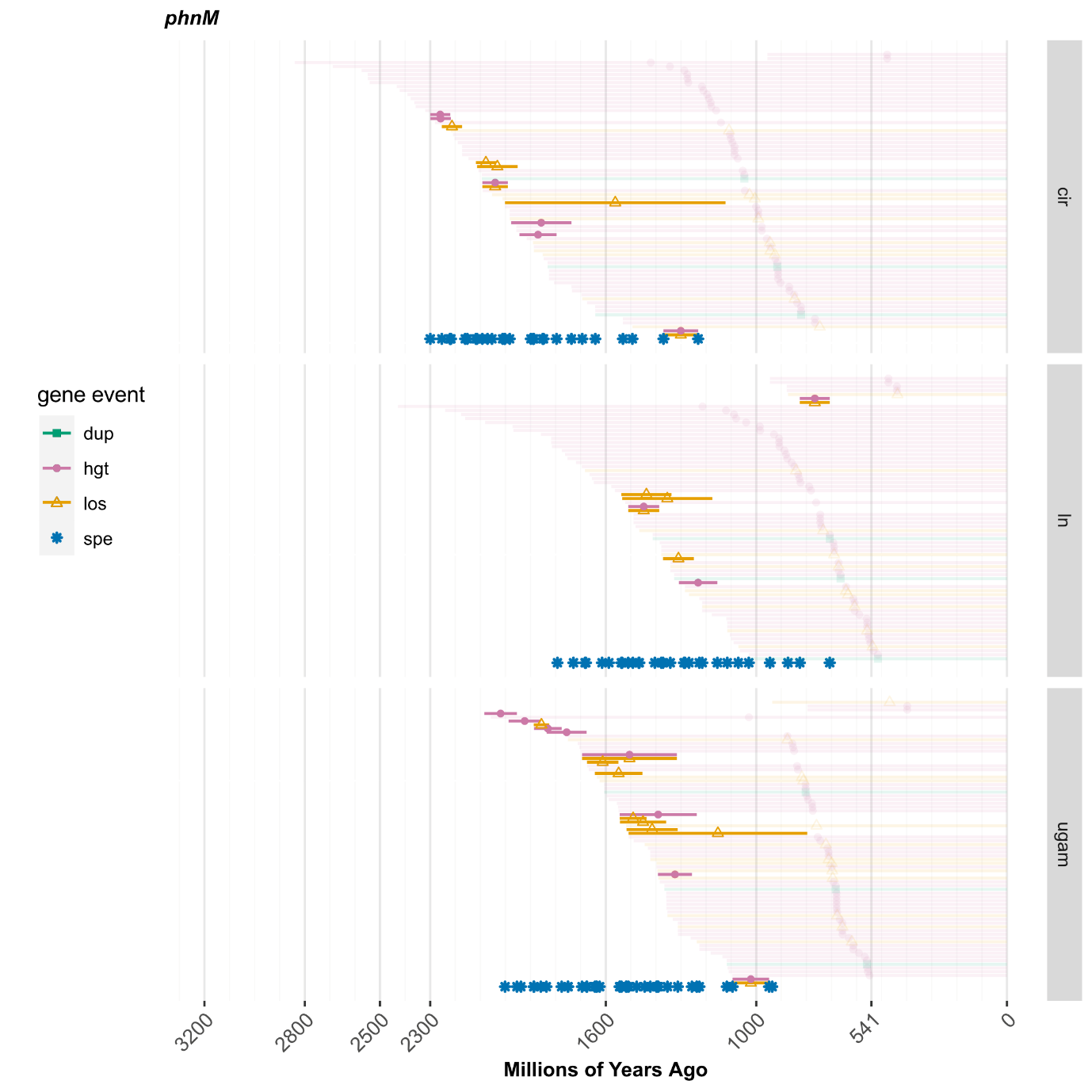

**Figure 1D: Uncertainty in estimating the origin of *phnM*.** Horizontal lines represent the lengths of branches where gene duplications (green), horizontal gene transfers (pink) and losses (orange) are predicted to have occurred. The midpoint of each branch is marked with shapes of the same colour, representing duplications (filled squares), horizontal gene transfers (filled circles) and losses (empty triangles). Darkness indicates whether the event occurred on an internal (dark colour) or terminal (faded) branch of the tree of life. Gene speciations (blue asterisks) are not associated with branch lengths because they occur on internal nodes of the tree. Results found using three different clock models are shown (cir: Cox-Ingersoll-Ross, ln: lognormal, ugam: uncorrelated gamma multipliers).

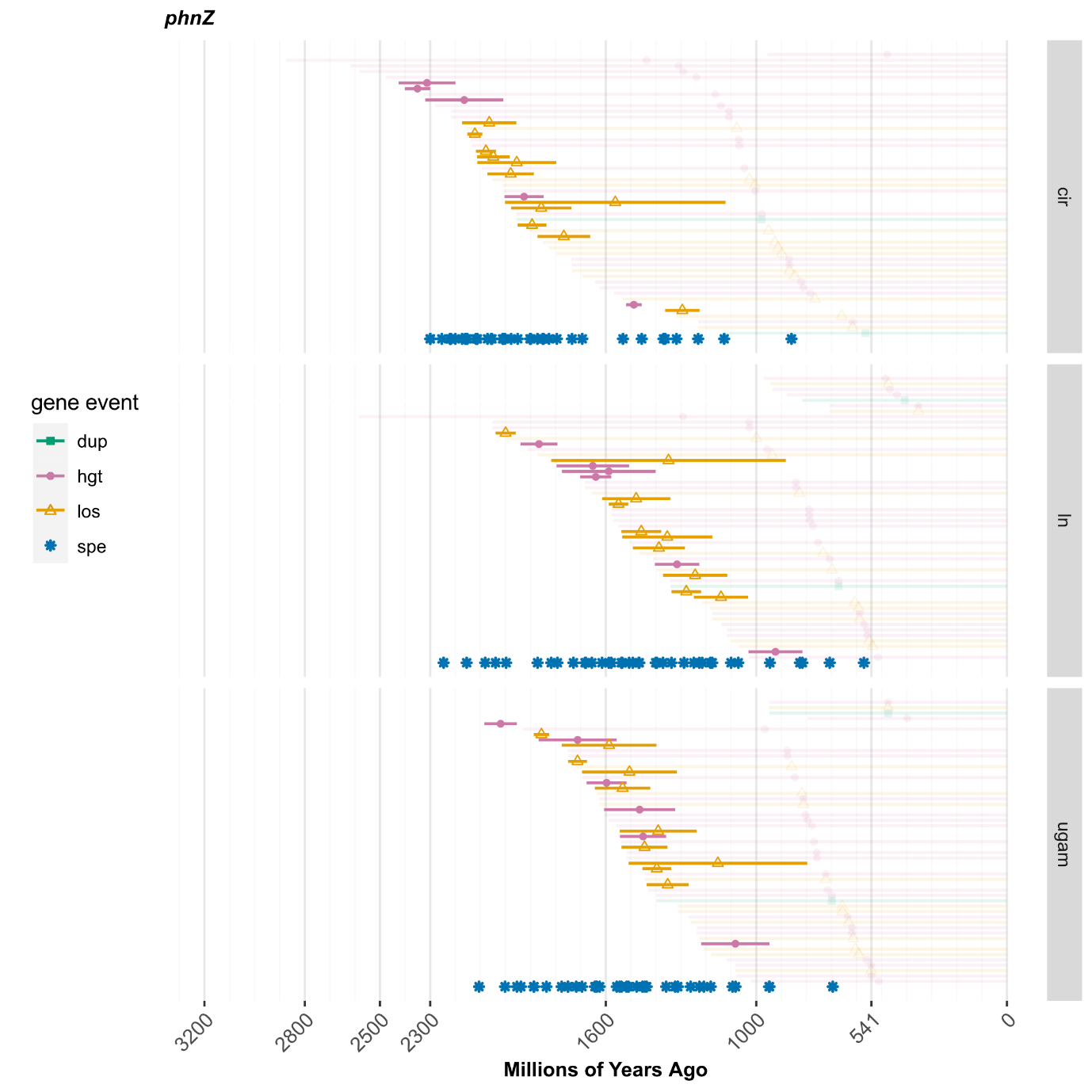

**Figure 1E: Uncertainty in estimating the origin of *phnZ*.** Horizontal lines represent the lengths of branches where gene duplications (green), horizontal gene transfers (pink) and losses (orange) are predicted to have occurred. The midpoint of each branch is marked with shapes of the same colour, representing duplications (filled squares), horizontal gene transfers (filled circles) and losses (empty triangles). Darkness indicates whether the event occurred on an internal (dark colour) or terminal (faded) branch of the tree of life. Gene speciations (blue asterisks) are not associated with branch lengths because they occur on internal nodes of the tree. Results found using three different clock models are shown (cir: Cox-Ingersoll-Ross, ln: lognormal, ugam: uncorrelated gamma multipliers).

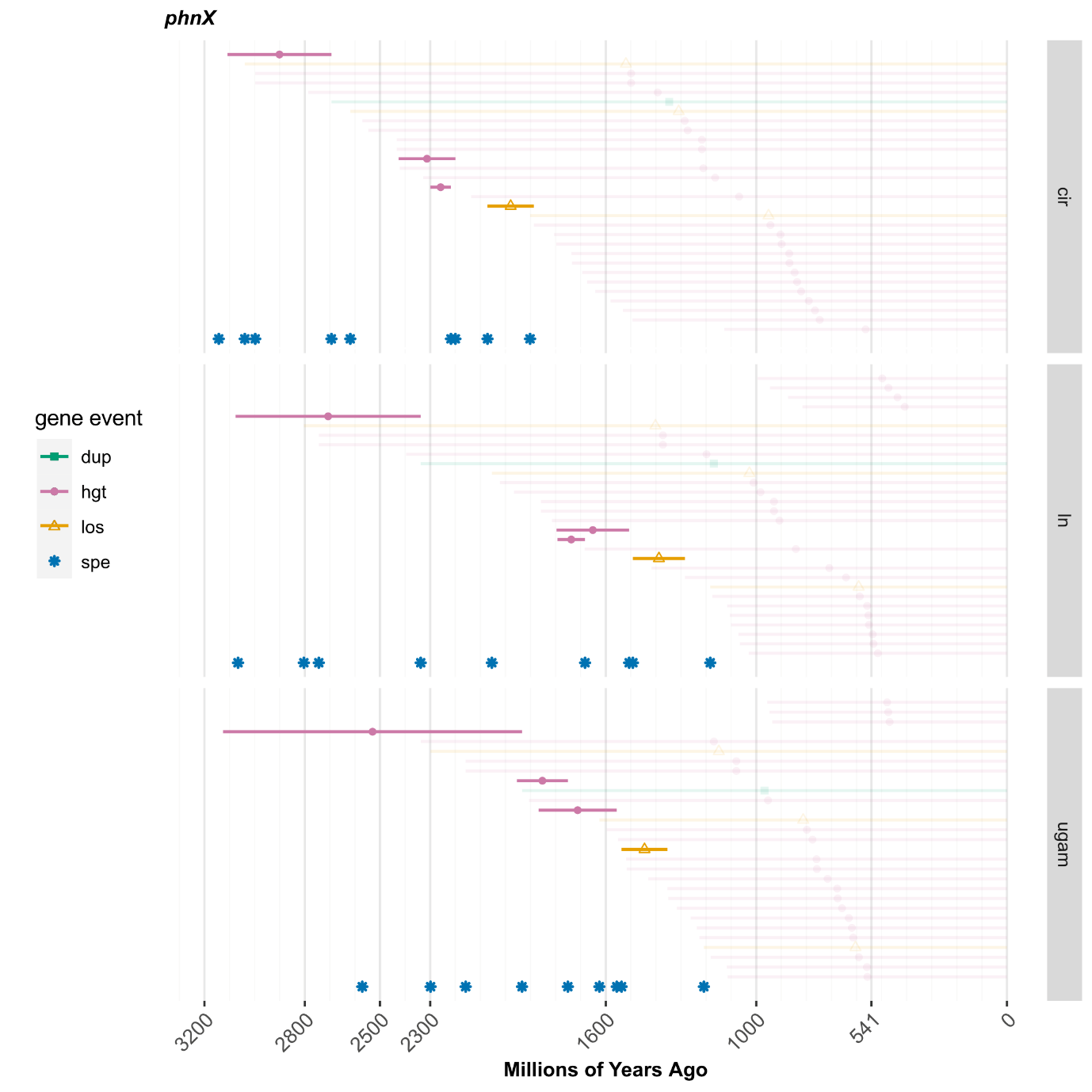

**Figure 1F: Uncertainty in estimating the origin of *phnX*.** Horizontal lines represent the lengths of branches where gene duplications (green), horizontal gene transfers (pink) and losses (orange) are predicted to have occurred. The midpoint of each branch is marked with shapes of the same colour, representing duplications (filled squares), horizontal gene transfers (filled circles) and losses (empty triangles). Darkness indicates whether the event occurred on an internal (dark colour) or terminal (faded) branch of the tree of life. Gene speciations (blue asterisks) are not associated with branch lengths because they occur on internal nodes of the tree. Results found using three different clock models are shown (cir: Cox-Ingersoll-Ross, ln: lognormal, ugam: uncorrelated gamma multipliers).

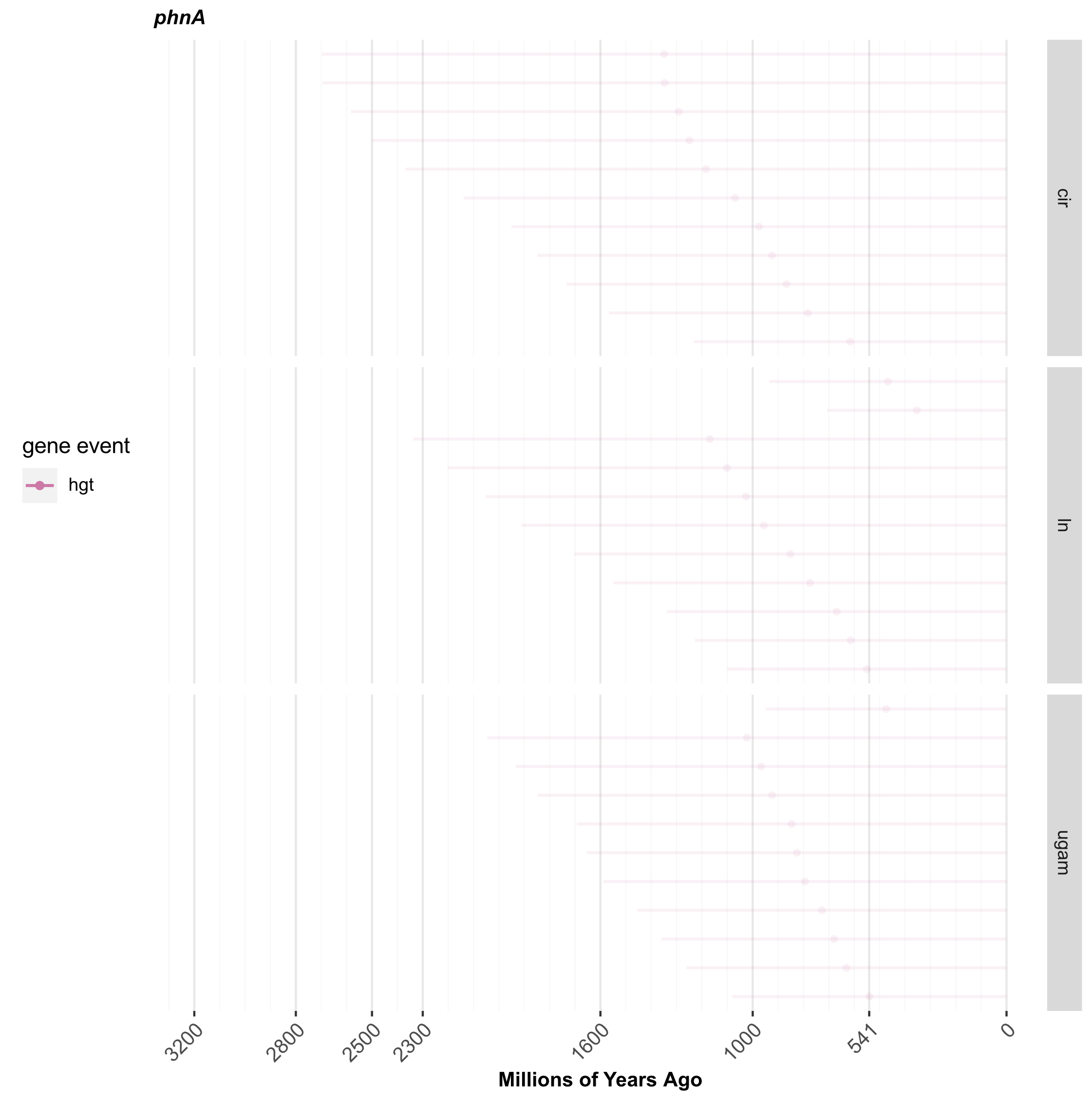

**Figure 1G: Uncertainty in estimating the origin of *phnA*.** Horizontal lines represent the lengths of branches where horizontal gene transfers (pink) are predicted to have occurred on internal branches of the tree of life. The midpoint of each branch is marked with filled circles of the same colour. Results found using three different clock models are shown (cir: Cox-Ingersoll-Ross, ln: lognormal, ugam: uncorrelated gamma multipliers).

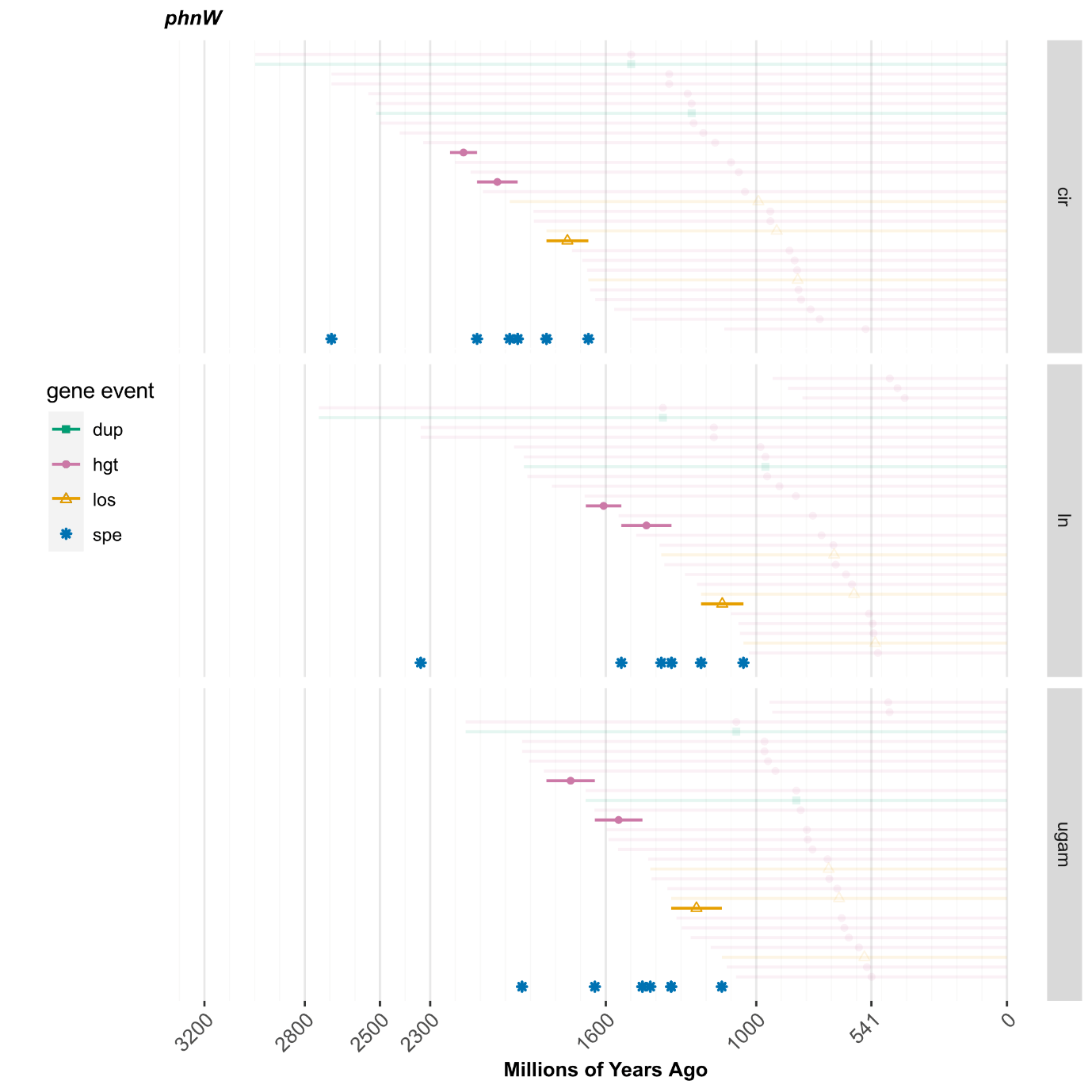

**Figure 1H: Uncertainty in estimating the origin of *phnW*.** Horizontal lines represent the lengths of branches where gene duplications (green), horizontal gene transfers (pink) and losses (orange) are predicted to have occurred. The midpoint of each branch is marked with shapes of the same colour, representing duplications (filled squares), horizontal gene transfers (filled circles) and losses (empty triangles). Darkness indicates whether the event occurred on an internal (dark colour) or terminal (faded) branch of the tree of life. Gene speciations (blue asterisks) are not associated with branch lengths because they occur on internal nodes of the tree. Results found using three different clock models are shown (cir: Cox-Ingersoll-Ross, ln: lognormal, ugam: uncorrelated gamma multipliers).

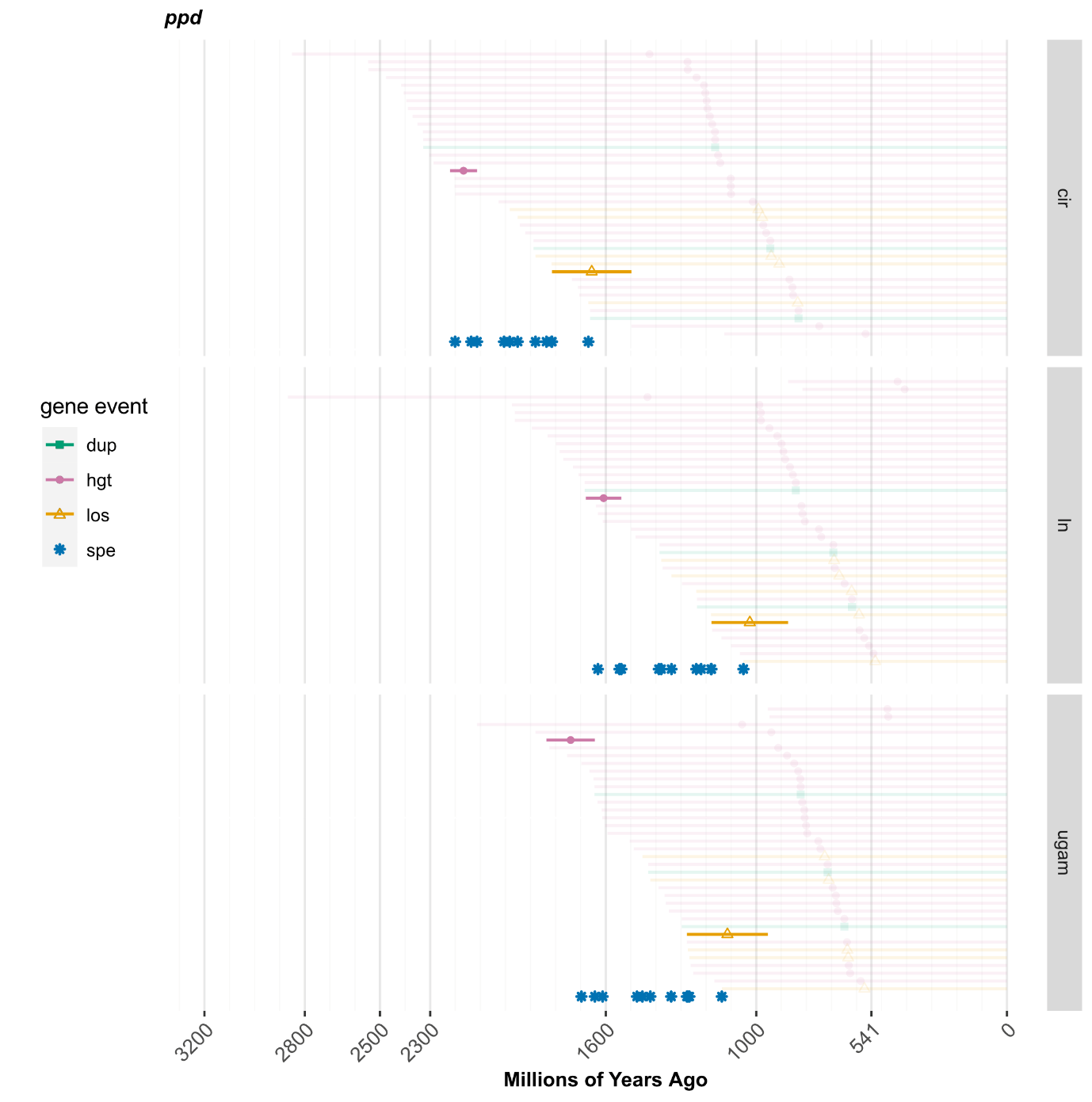

**Figure 1I: Uncertainty in estimating the origin of *ppd*.** Horizontal lines represent the lengths of branches where gene duplications (green), horizontal gene transfers (pink) and losses (orange) are predicted to have occurred. The midpoint of each branch is marked with shapes of the same colour, representing duplications (filled squares), horizontal gene transfers (filled circles) and losses (empty triangles). Darkness indicates whether the event occurred on an internal (dark colour) or terminal (faded) branch of the tree of life. Gene speciations (blue asterisks) are not associated with branch lengths because they occur on internal nodes of the tree. Results found using three different clock models are shown (cir: Cox-Ingersoll-Ross, ln: lognormal, ugam: uncorrelated gamma multipliers).

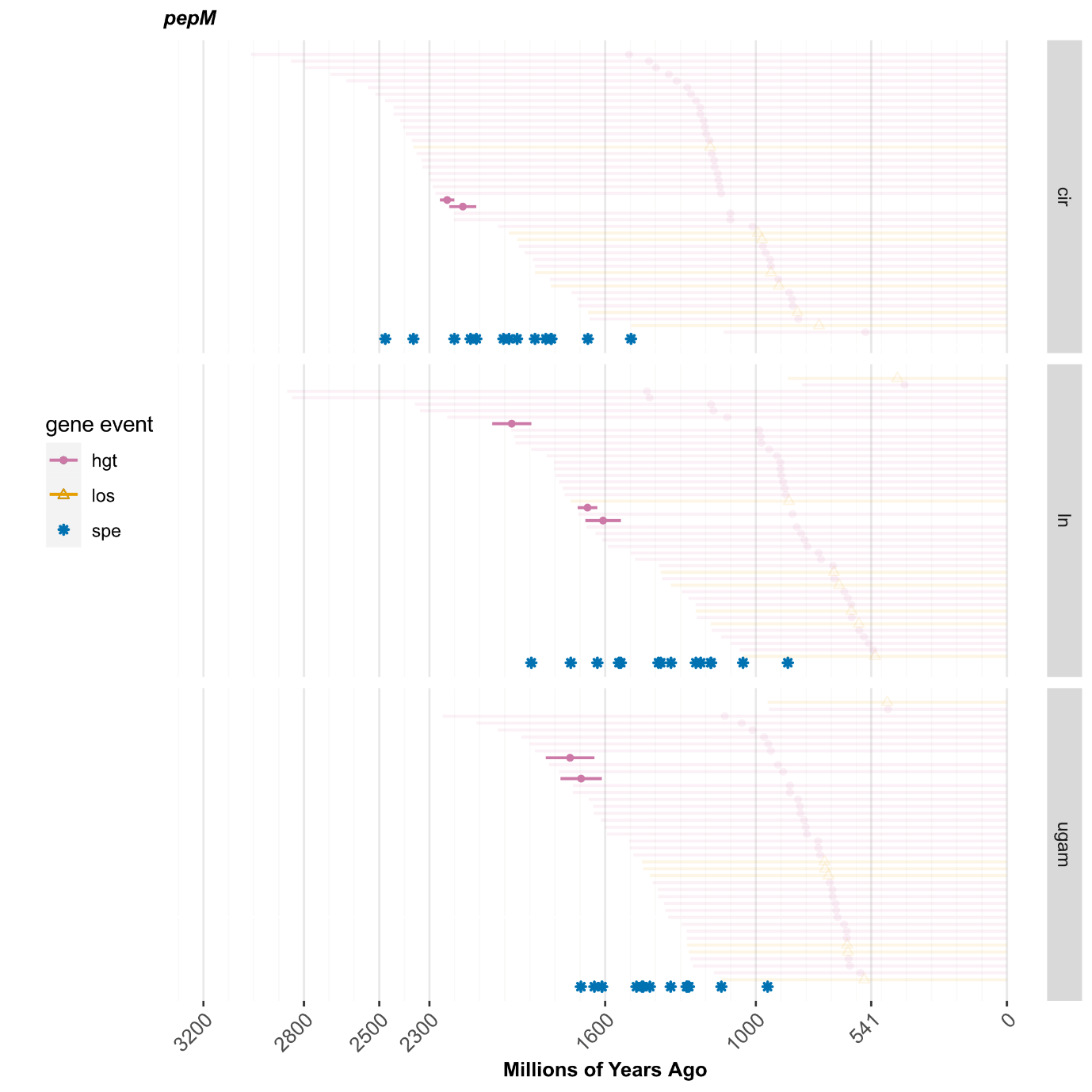

**Figure 1J: Uncertainty in estimating the origin of *pepM*.** Horizontal lines represent the lengths of branches where gene duplications (green), horizontal gene transfers (pink) and losses (orange) are predicted to have occurred. The midpoint of each branch is marked with shapes of the same colour, representing duplications (filled squares), horizontal gene transfers (filled circles) and losses (empty triangles). Darkness indicates whether the event occurred on an internal (dark colour) or terminal (faded) branch of the tree of life. Gene speciations (blue asterisks) are not associated with branch lengths because they occur on internal nodes of the tree. Results found using three different clock models are shown (cir: Cox-Ingersoll-Ross, ln: lognormal, ugam: uncorrelated gamma multipliers).

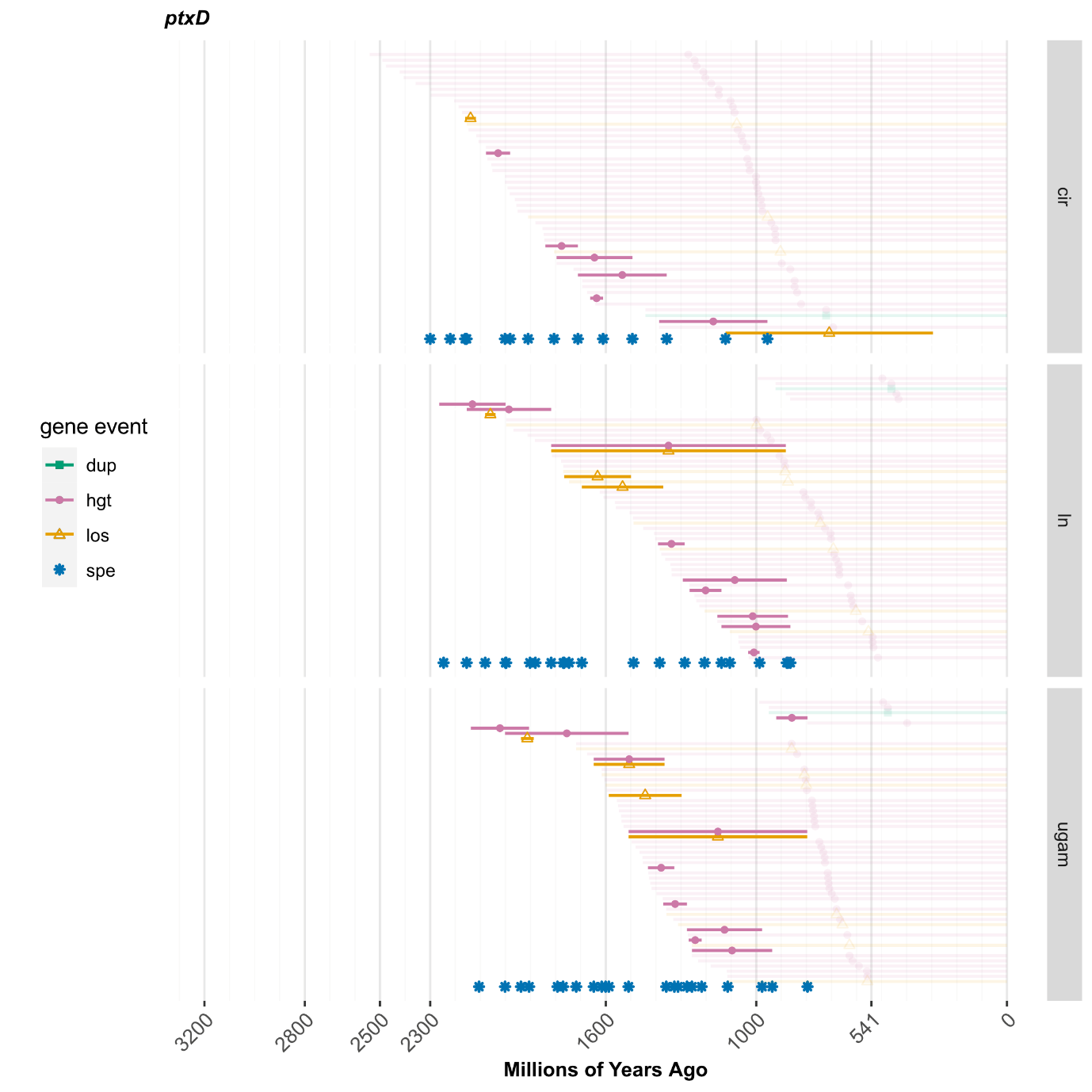

**Figure 1K: Uncertainty in estimating the origin of *ptxD*.** Horizontal lines represent the lengths of branches where gene duplications (green), horizontal gene transfers (pink) and losses (orange) are predicted to have occurred. The midpoint of each branch is marked with shapes of the same colour, representing duplications (filled squares), horizontal gene transfers (filled circles) and losses (empty triangles). Darkness indicates whether the event occurred on an internal (dark colour) or terminal (faded) branch of the tree of life. Gene speciations (blue asterisks) are not associated with branch lengths because they occur on internal nodes of the tree. Results found using three different clock models are shown (cir: Cox-Ingersoll-Ross, ln: lognormal, ugam: uncorrelated gamma multipliers).

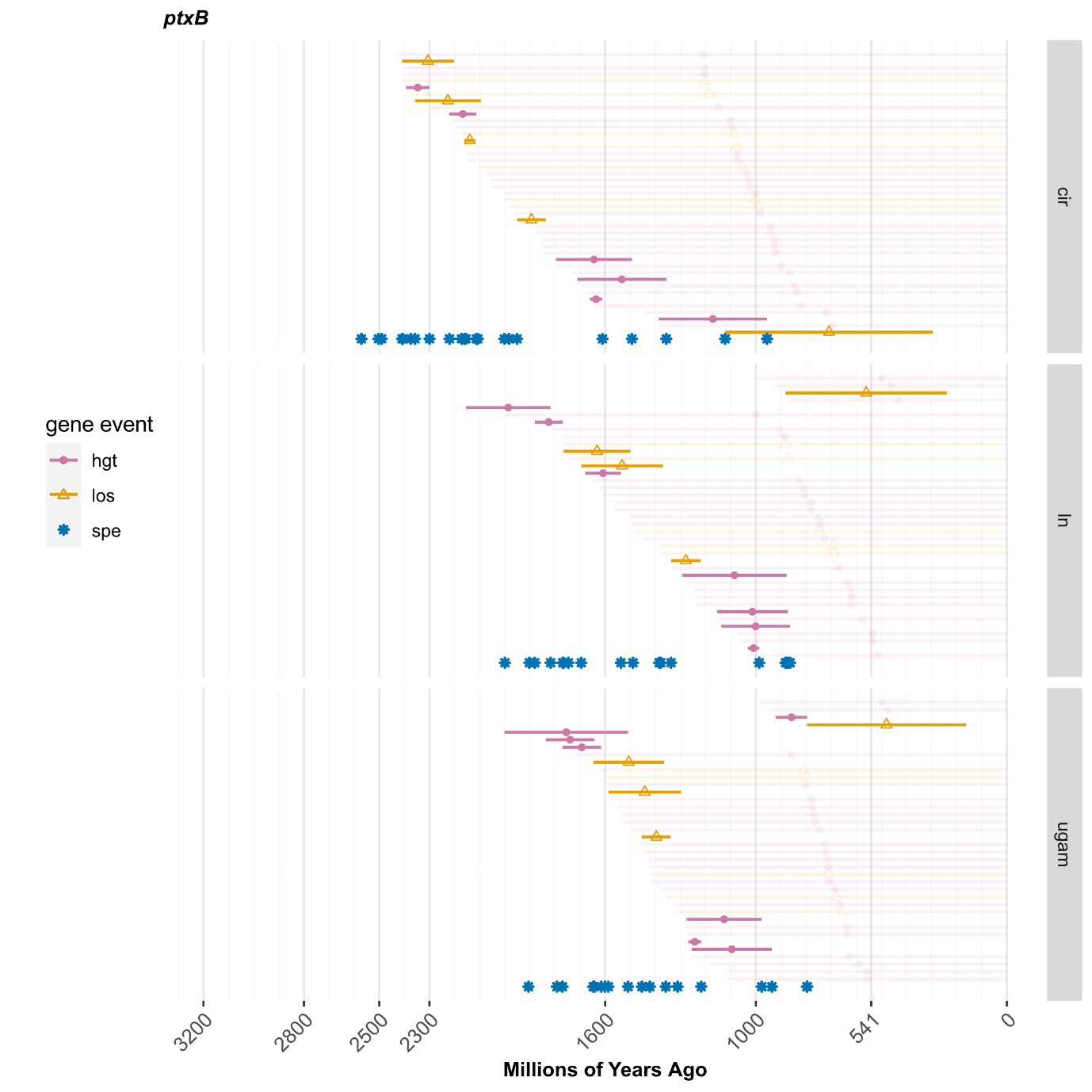

**Figure 1L: Uncertainty in estimating the origin of *ptxB*.** Horizontal lines represent the lengths of branches where gene duplications (green), horizontal gene transfers (pink) and losses (orange) are predicted to have occurred. The midpoint of each branch is marked with shapes of the same colour, representing duplications (filled squares), horizontal gene transfers (filled circles) and losses (empty triangles). Darkness indicates whether the event occurred on an internal (dark colour) or terminal (faded) branch of the tree of life. Gene speciations (blue asterisks) are not associated with branch lengths because they occur on internal nodes of the tree. Results found using three different clock models are shown (cir: Cox-Ingersoll-Ross, ln: lognormal, ugam: uncorrelated gamma multipliers).

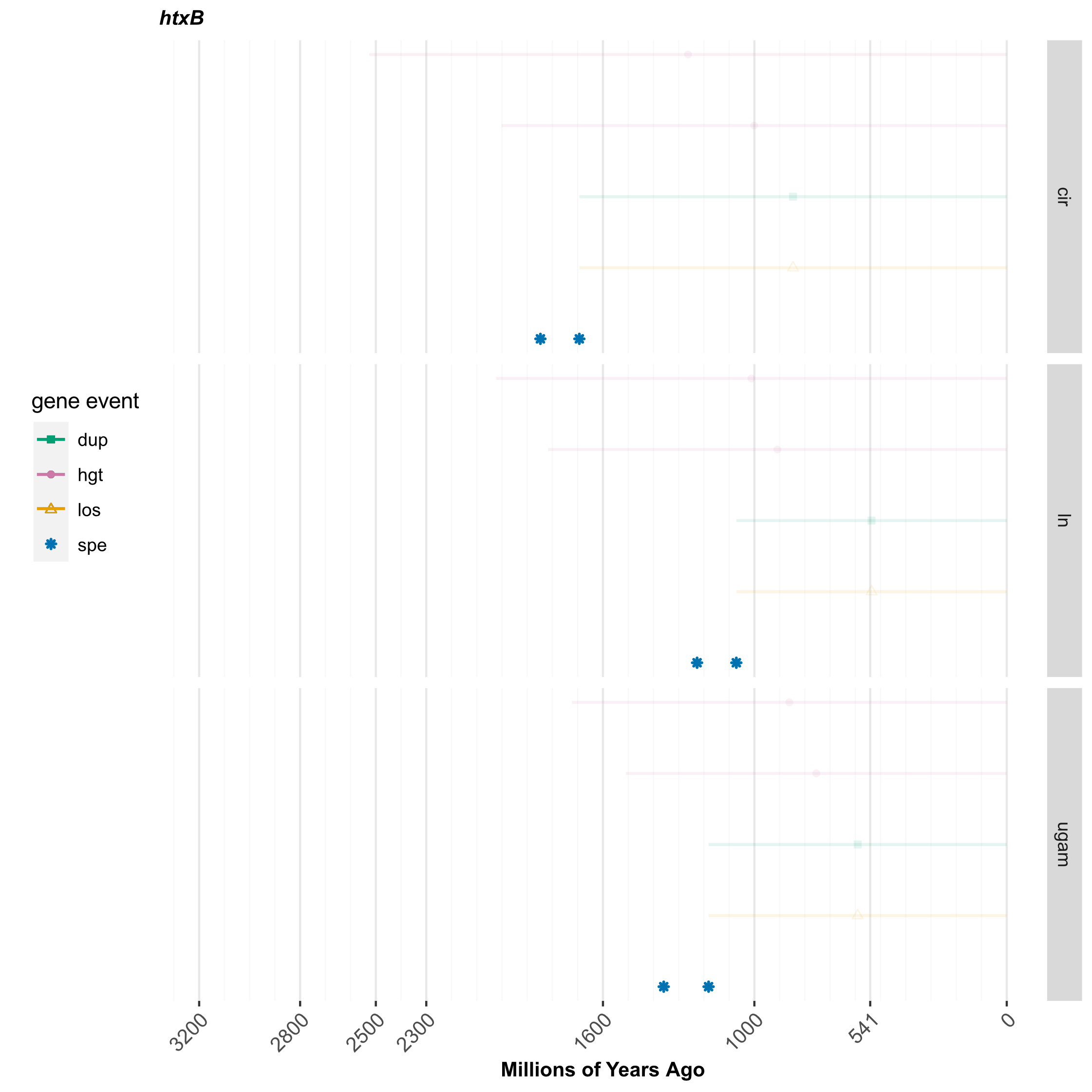

**Figure 1M: Uncertainty in estimating the origin of *htxB*.** Horizontal lines represent the lengths of branches where gene duplications (green), horizontal gene transfers (pink) and losses (orange) are predicted to have occurred. The midpoint of each branch is marked with shapes of the same colour, representing duplications (filled squares), horizontal gene transfers (filled circles) and losses (empty triangles). Darkness indicates whether the event occurred on an internal (dark colour) or terminal (faded) branch of the tree of life. Gene speciations (blue asterisks) are not associated with branch lengths because they occur on internal nodes of the tree. Results found using three different clock models are shown (cir: Cox-Ingersoll-Ross, ln: lognormal, ugam: uncorrelated gamma multipliers).

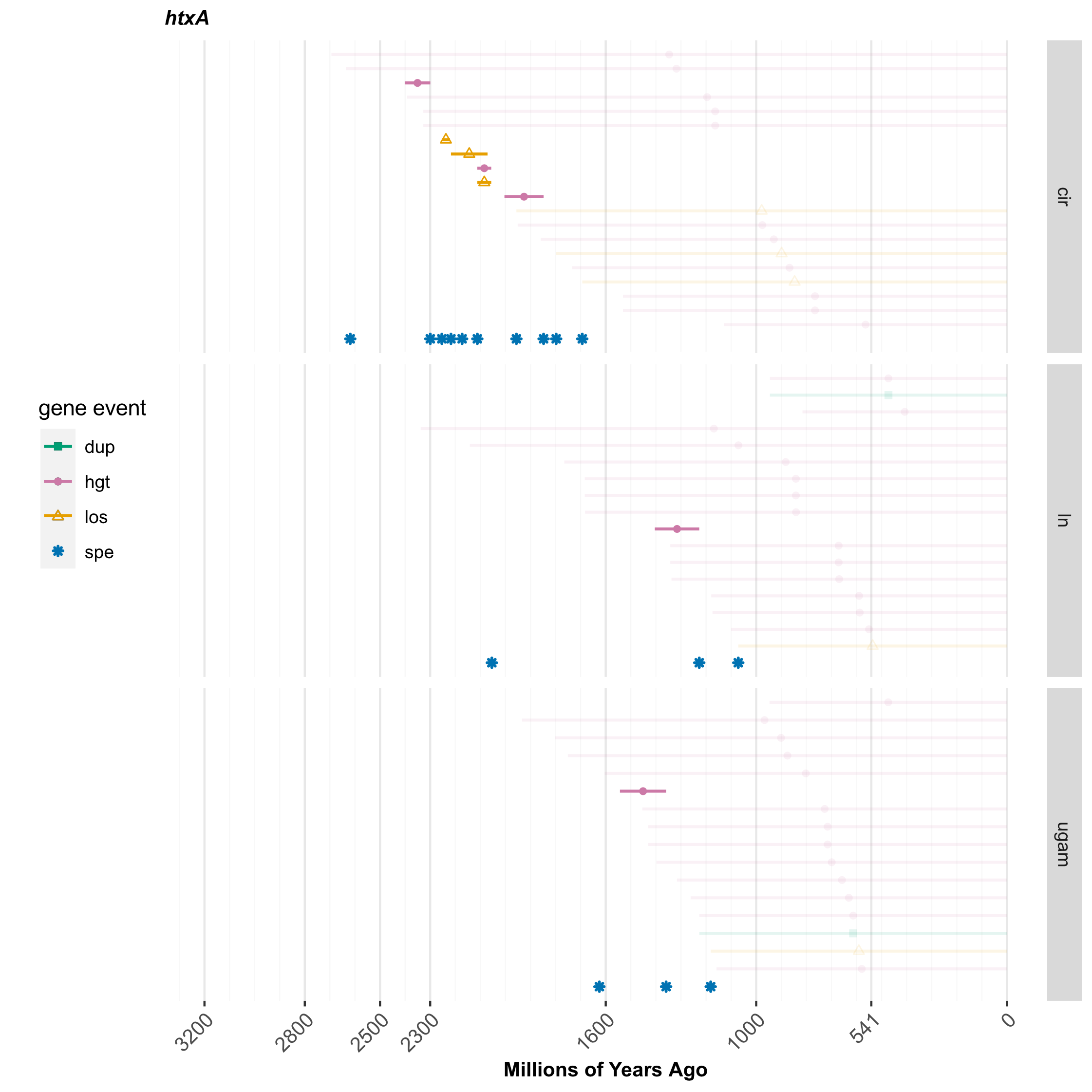

**Figure 1N: Uncertainty in estimating the origin of *htxA*.** Horizontal lines represent the lengths of branches where gene duplications (green), horizontal gene transfers (pink) and losses (orange) are predicted to have occurred. The midpoint of each branch is marked with shapes of the same colour, representing duplications (filled squares), horizontal gene transfers (filled circles) and losses (empty triangles). Darkness indicates whether the event occurred on an internal (dark colour) or terminal (faded) branch of the tree of life. Gene speciations (blue asterisks) are not associated with branch lengths because they occur on internal nodes of the tree. Results found using three different clock models are shown (cir: Cox-Ingersoll-Ross, ln: lognormal, ugam: uncorrelated gamma multipliers).

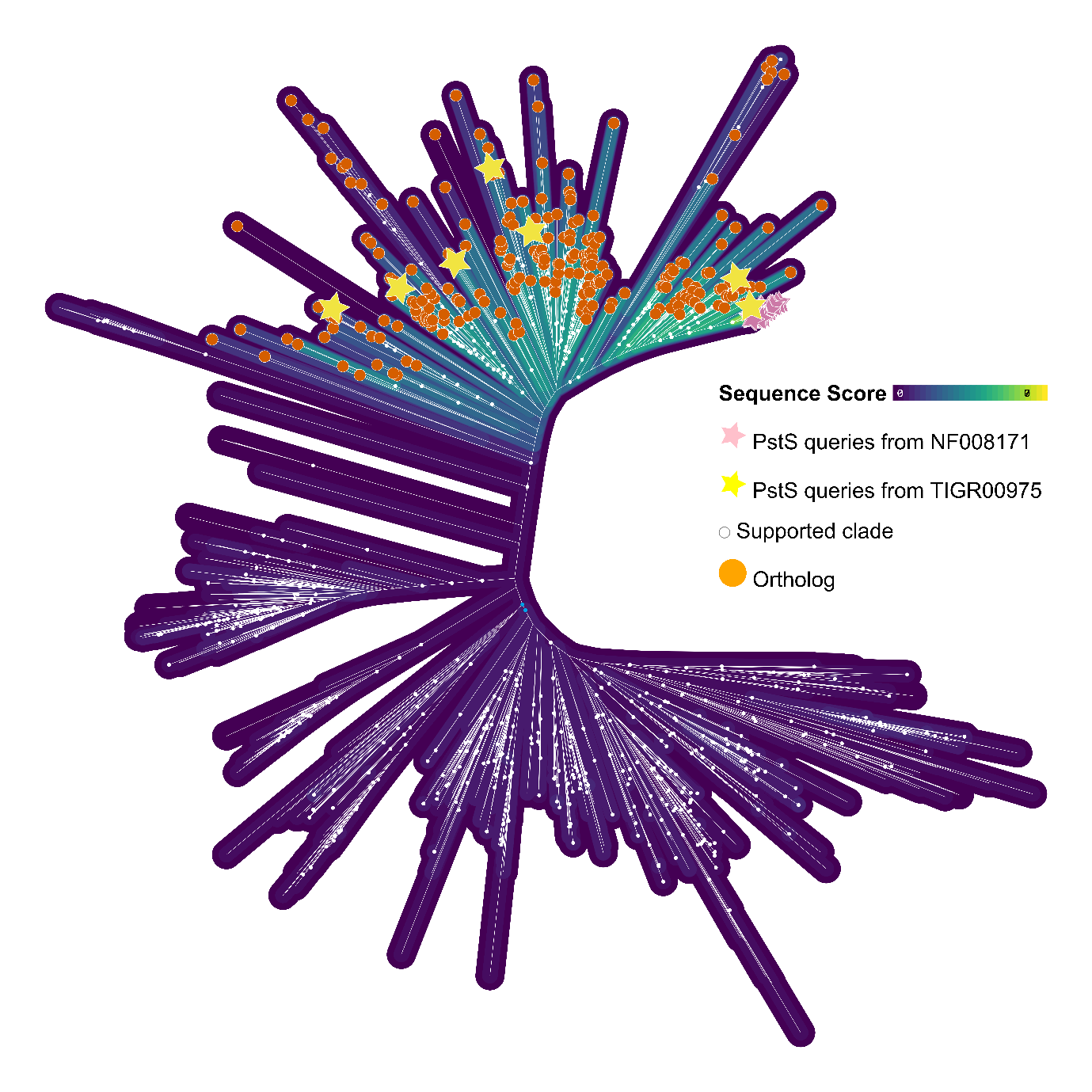

**Figure 2A: Identification of *pstS* orthologs.** Maximum-likelihood phylogeny of *pstS* homologs, with bitscores represented by coloured outlines surrounding each branch. Orthologs (orange circles) were identified and selected for further analyses based on their bitscore and relationship to query sequences (yellow stars and pink stars) from the respective HMM profiles. White circles indicate ultrafast bootstrap support values => 95.

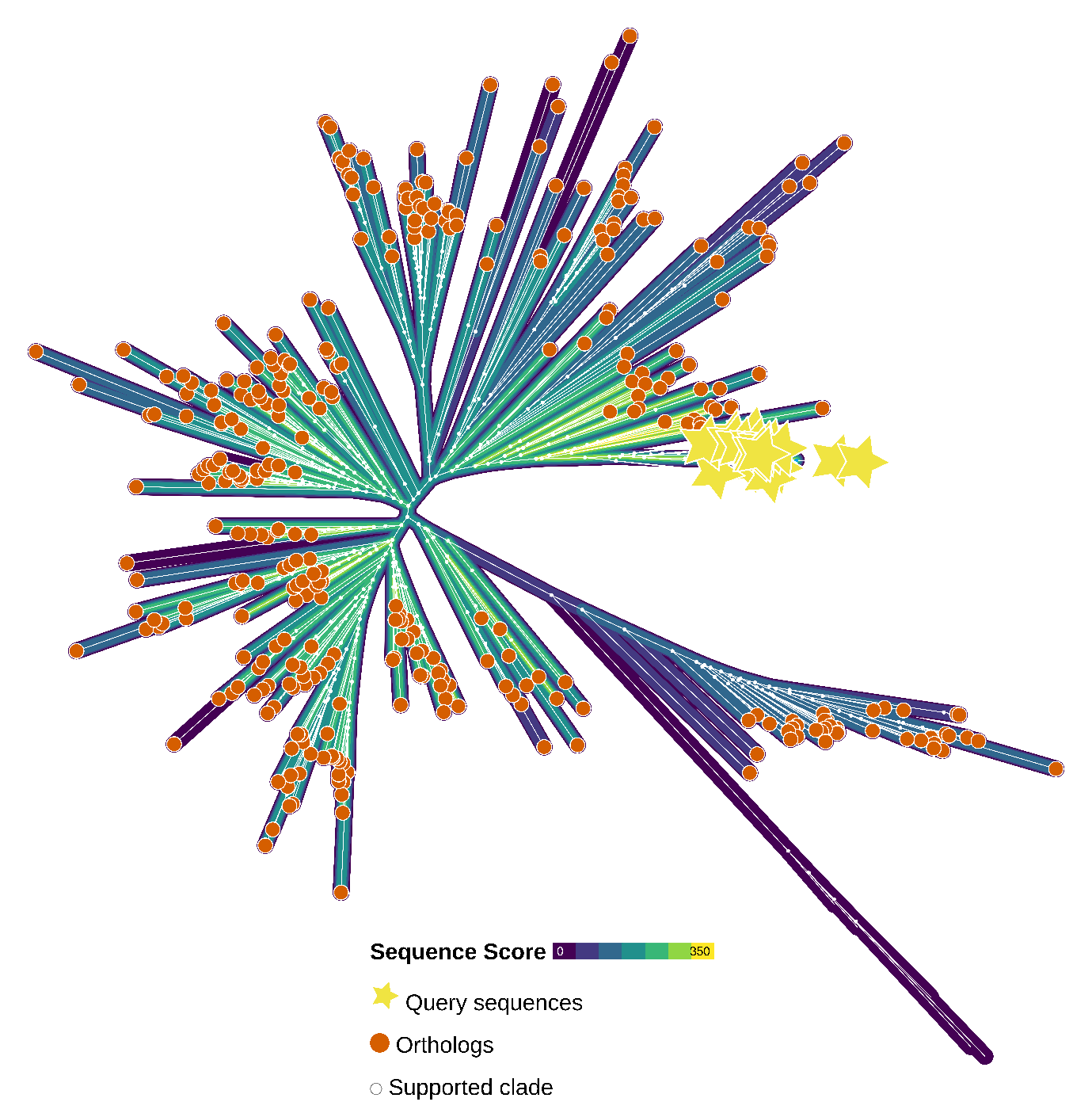

**Figure 2B: Identification of *pnas* orthologs.** Maximum-likelihood phylogeny of *pnas* homologs, with bitscores represented by coloured outlines surrounding each branch. Orthologs (orange circles) were identified and selected for further analyses based on their bitscore and relationship to query sequences (yellow stars) from the respective HMM profiles. White circles indicate ultrafast bootstrap support values => 95.

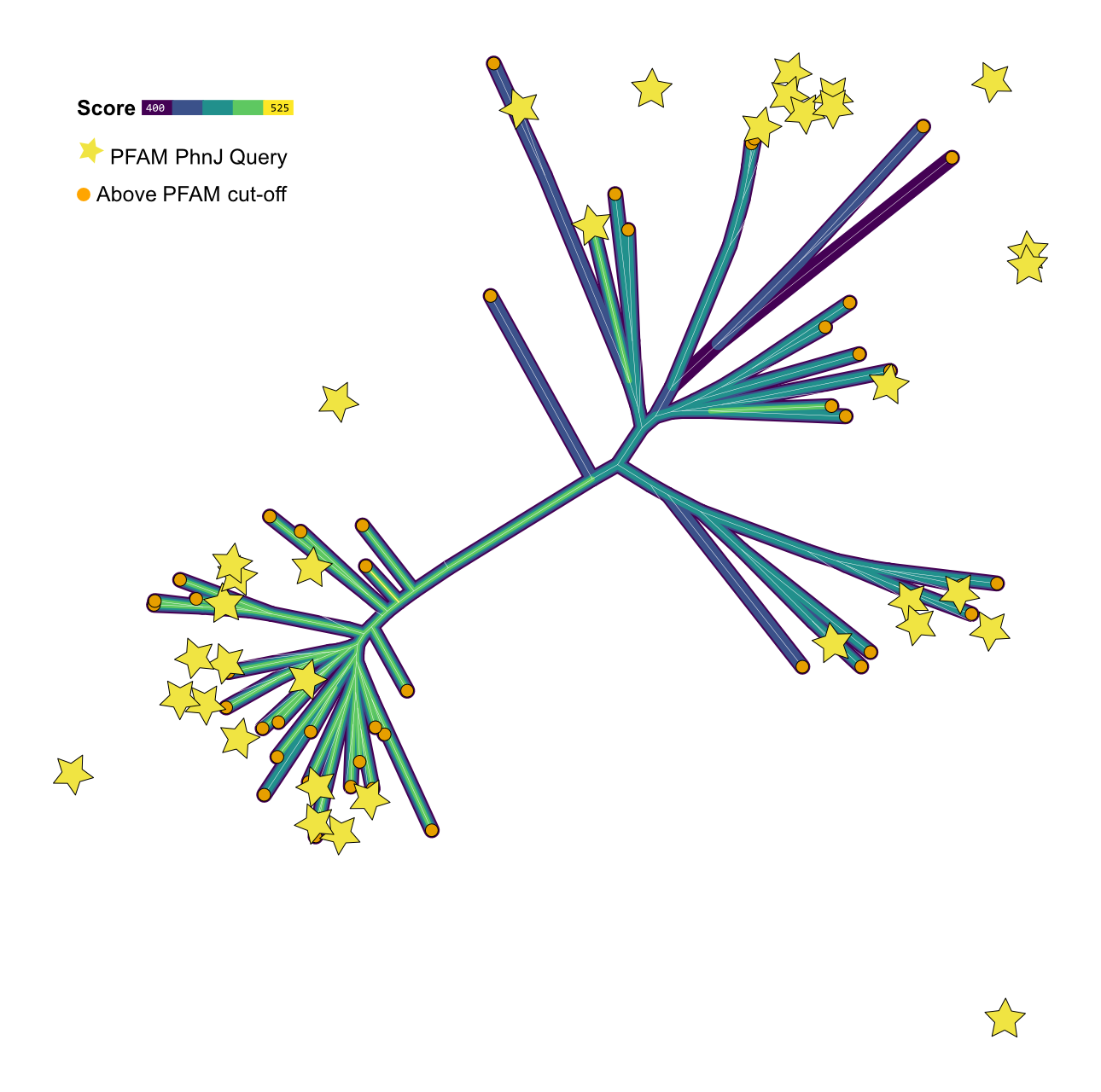

**Figure 2C: Identification of *phnJ* orthologs.** Maximum-likelihood phylogeny of *phnJ* homologs, with bitscores represented by coloured outlines surrounding each branch. Orthologs (orange circles) were identified and selected for further analyses based on their bitscore and relationship to query sequences (yellow stars) from the respective HMM profiles. White circles indicate ultrafast bootstrap support values => 95.

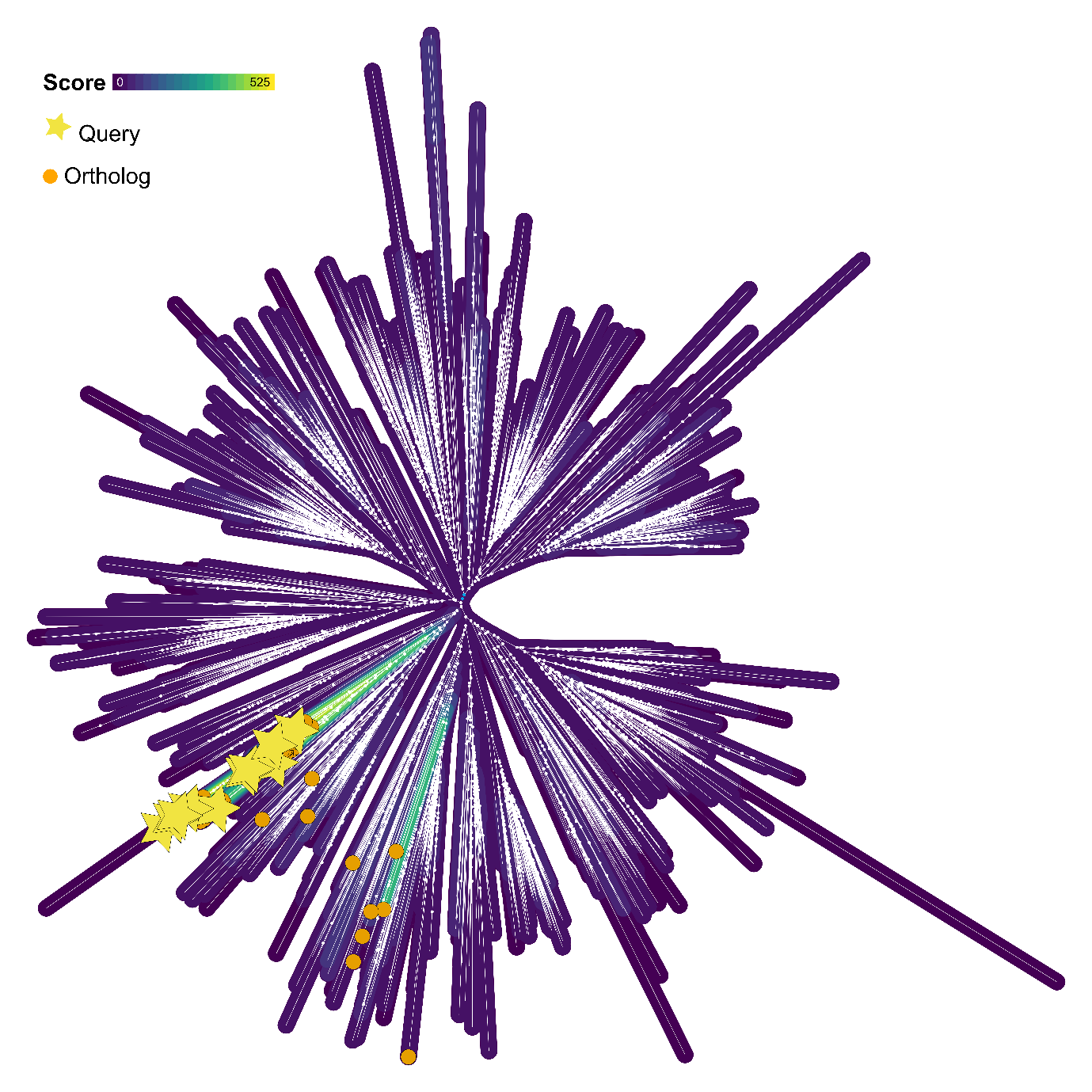

**Figure 2D: Identification of *phnM* orthologs.** Maximum-likelihood phylogeny of *phnM* homologs, with bitscores represented by coloured outlines surrounding each branch. Orthologs (orange circles) were identified and selected for further analyses based on their bitscore and relationship to query sequences (yellow stars) from the respective HMM profiles. White circles indicate ultrafast bootstrap support values => 95.

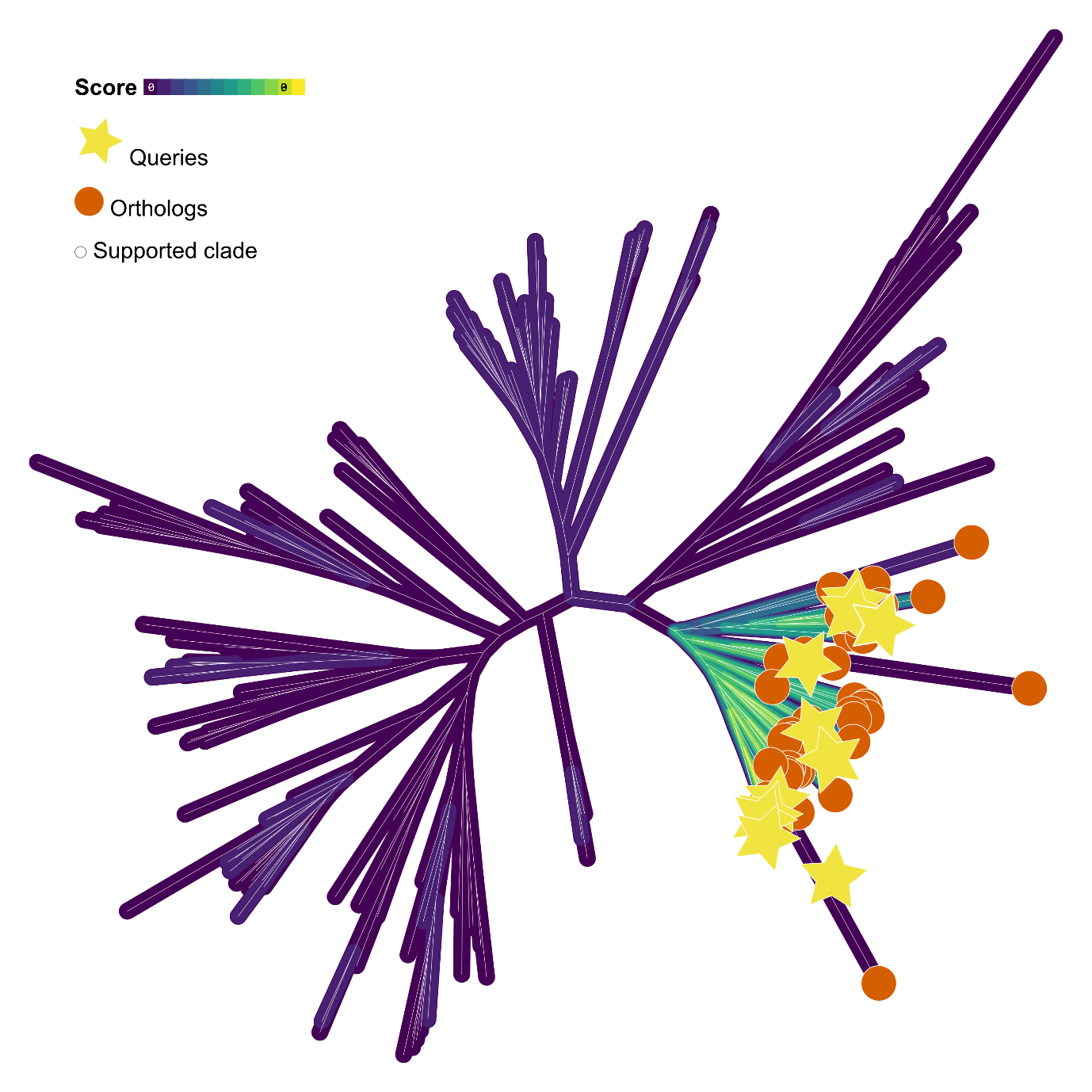

**Figure 2E: Identification of *phnZ* orthologs.** Maximum-likelihood phylogeny of *phnZ* homologs, with bitscores represented by coloured outlines surrounding each branch. Orthologs (orange circles) were identified and selected for further analyses based on their bitscore and relationship to query sequences (yellow stars) from the respective HMM profiles. White circles indicate ultrafast bootstrap support values => 95.

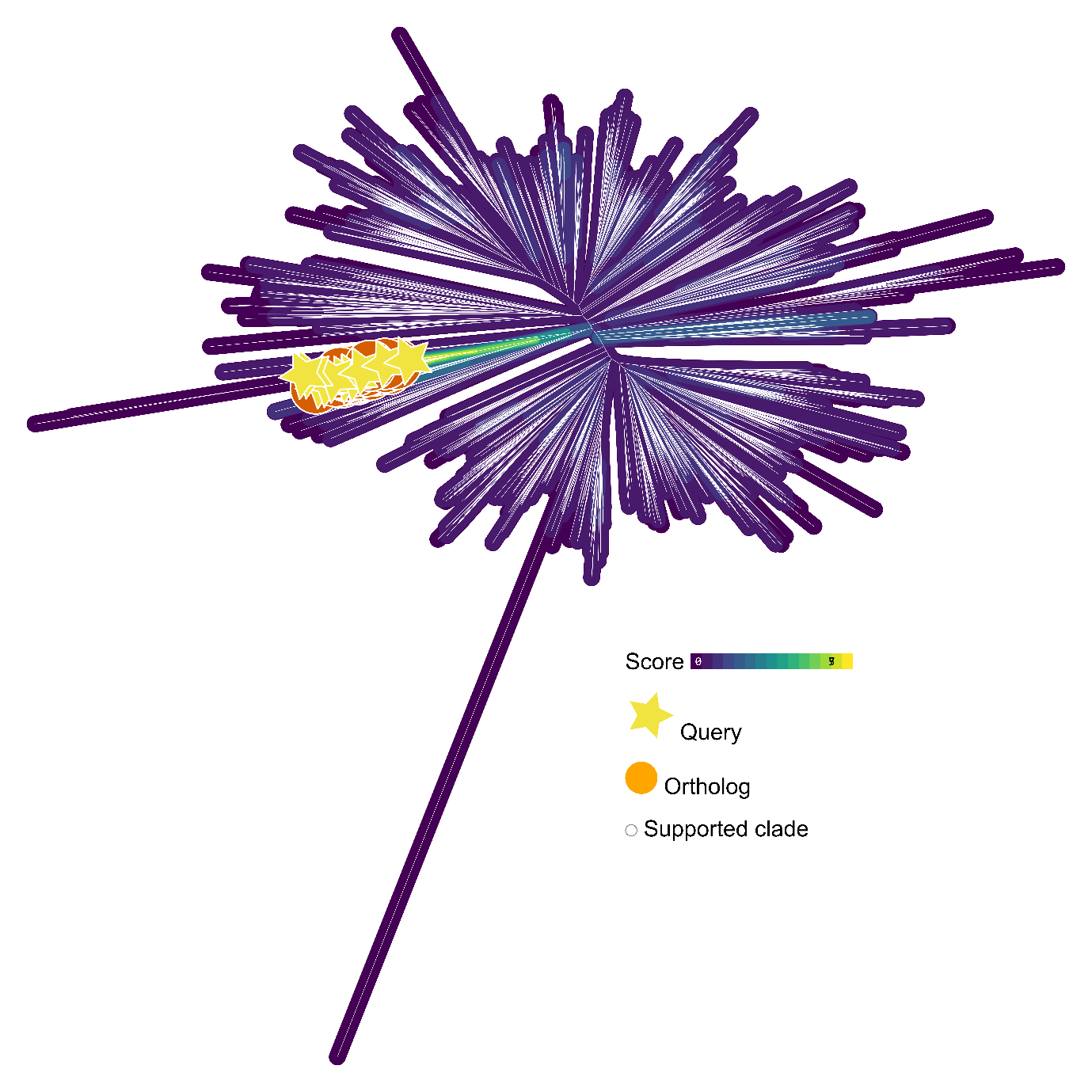

**Figure 2F: Identification of *phnX* orthologs.** Maximum-likelihood phylogeny of *phnX* homologs, with bitscores represented by coloured outlines surrounding each branch. Orthologs (orange circles) were identified and selected for further analyses based on their bitscore and relationship to query sequences (yellow stars) from the respective HMM profiles. White circles indicate ultrafast bootstrap support values => 95.

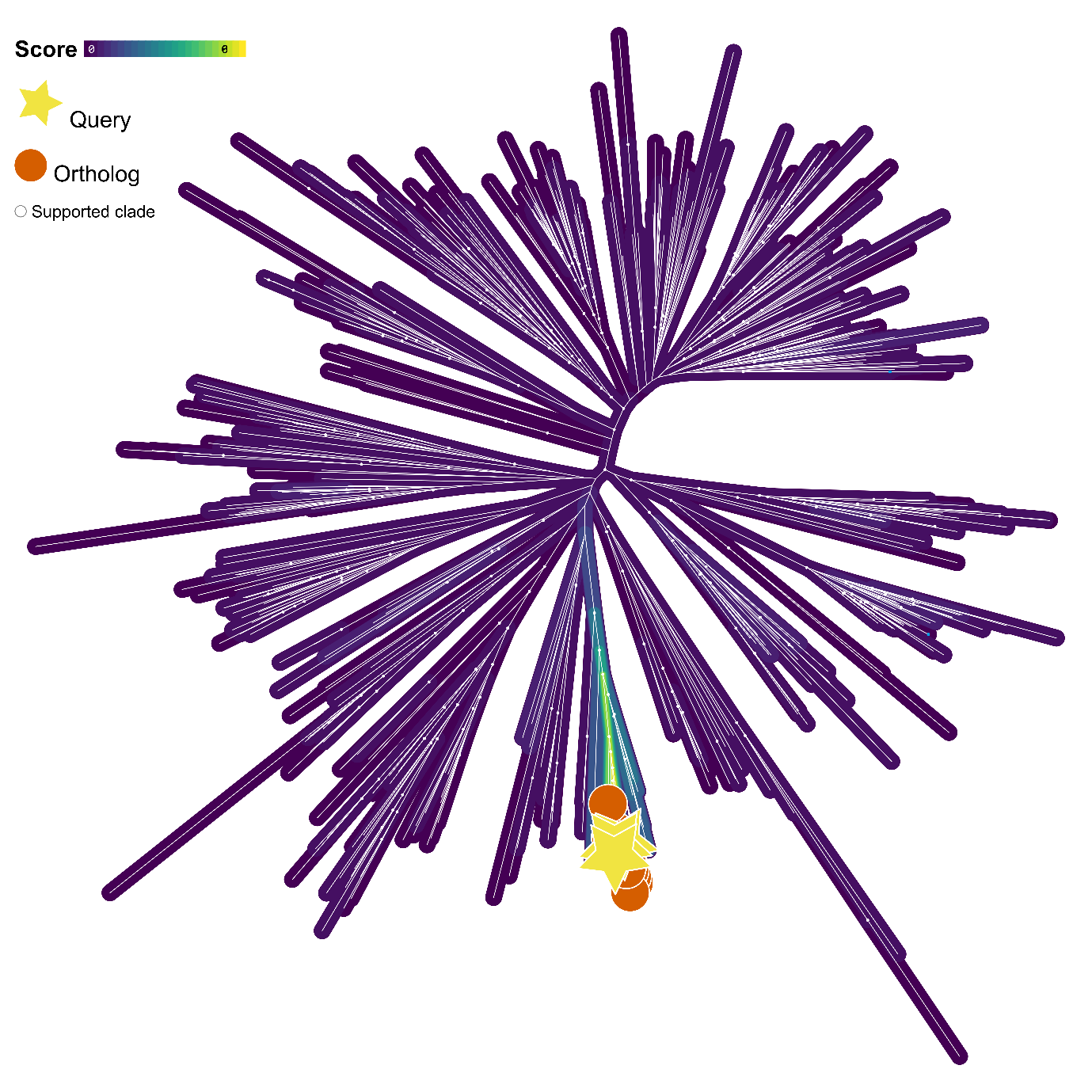

**Figure 2G: Identification of *phnA* orthologs.** Maximum-likelihood phylogeny of *phnA* homologs, with bitscores represented by coloured outlines surrounding each branch. Orthologs (orange circles) were identified and selected for further analyses based on their bitscore and relationship to query sequences (yellow stars) from the respective HMM profiles. White circles indicate ultrafast bootstrap support values => 95.

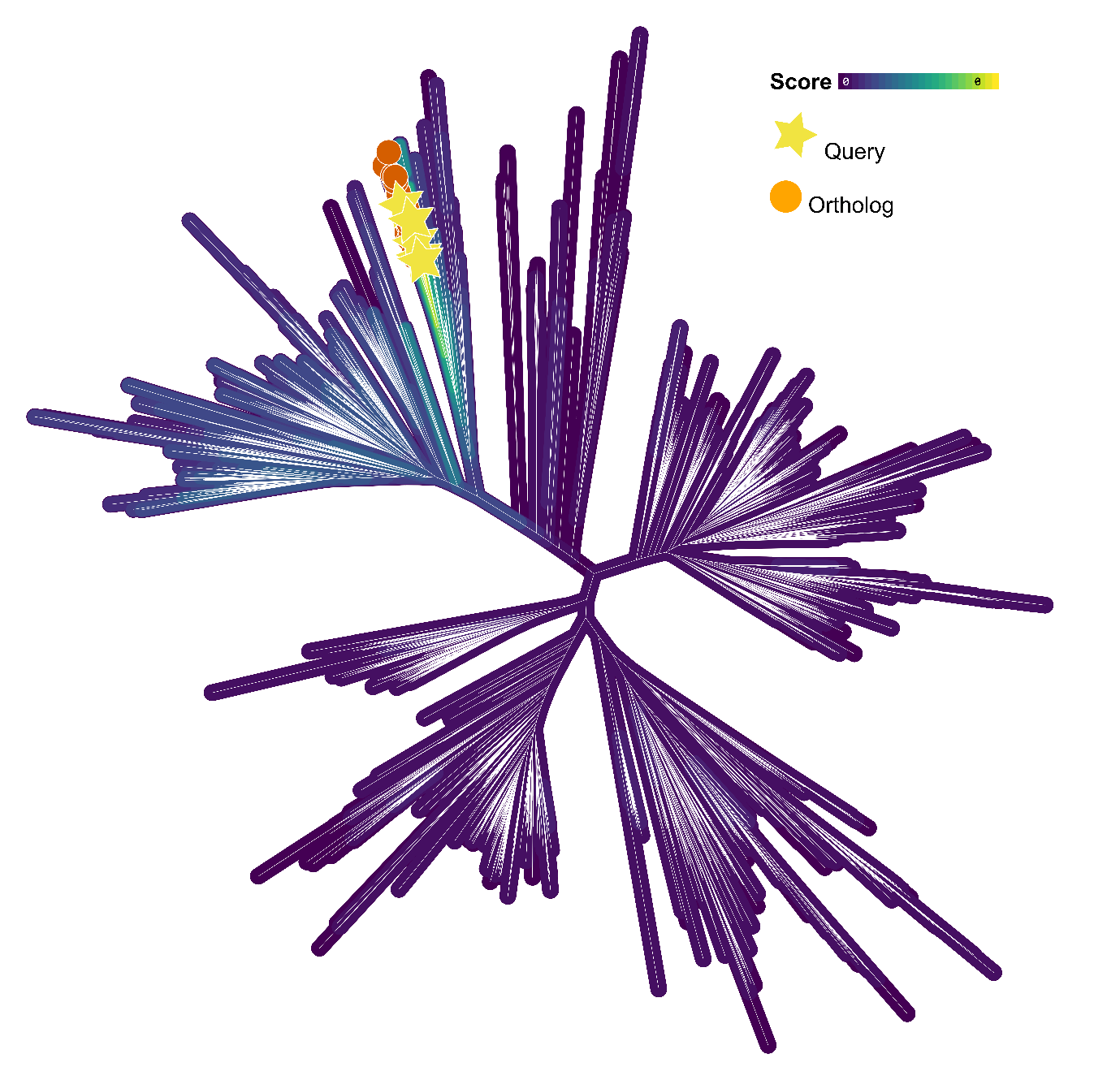

**Figure 2H: Identification of *phnW* orthologs.** Maximum-likelihood phylogeny of *phnW* homologs, with bitscores represented by coloured outlines surrounding each branch. Orthologs (orange circles) were identified and selected for further analyses based on their bitscore and relationship to query sequences (yellow stars) from the respective HMM profiles.

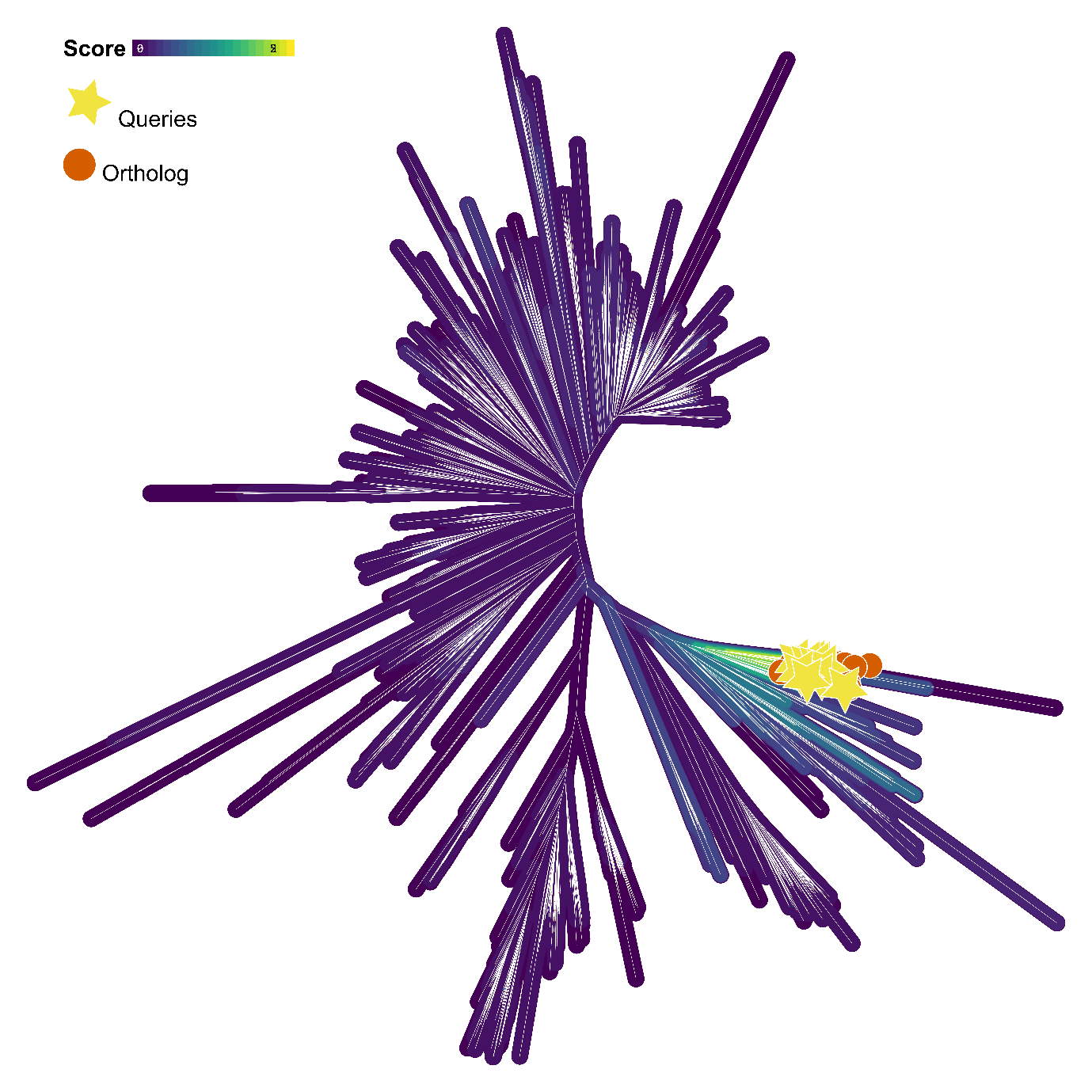

**Figure 2I: Identification of *ppd* orthologs.** Maximum-likelihood phylogeny of *ppd* homologs, with bitscores represented by coloured outlines surrounding each branch. Orthologs (orange circles) were identified and selected for further analyses based on their bitscore and relationship to query sequences (yellow stars) from the respective HMM profiles.

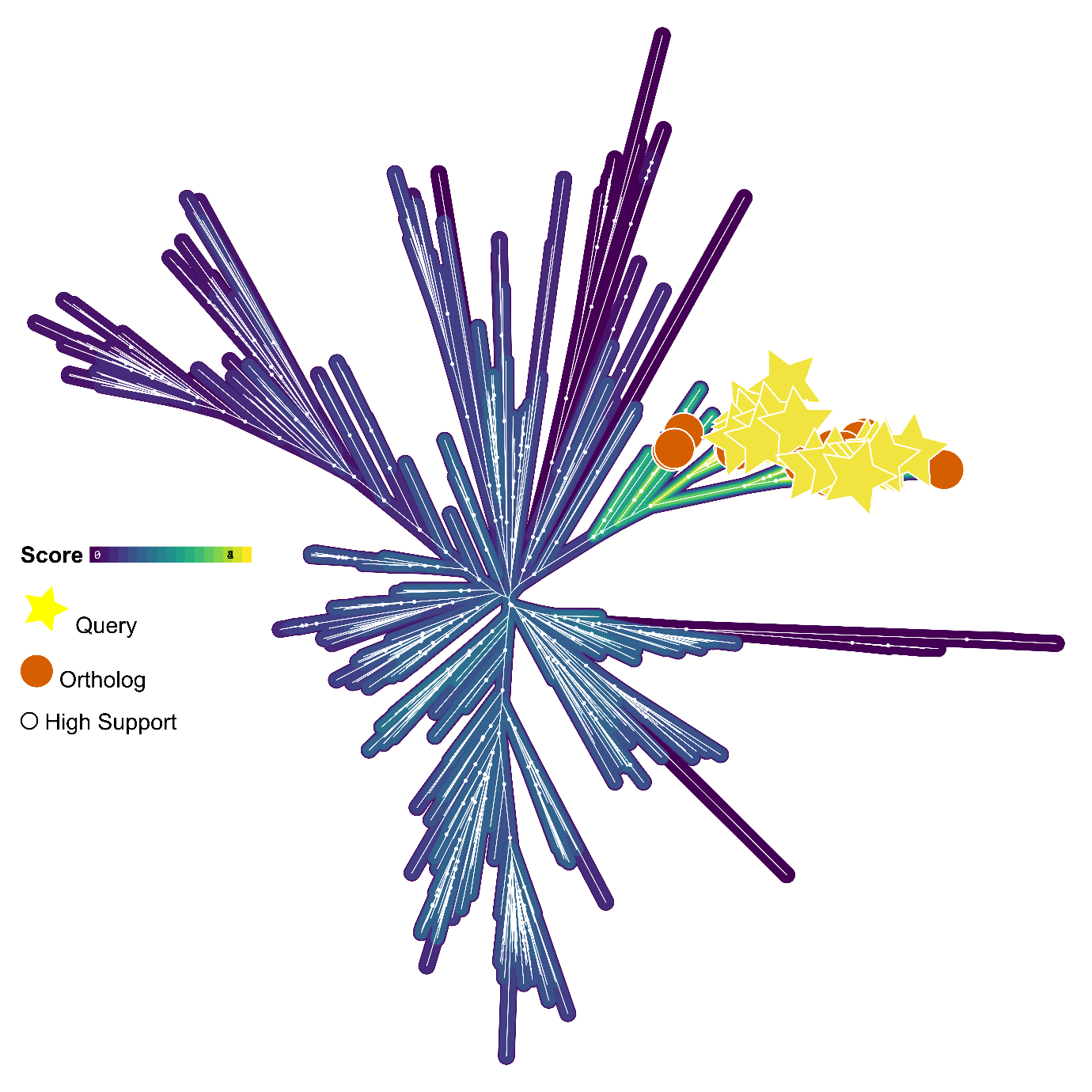

**Figure 2J: Identification of *pepM* orthologs.** Maximum-likelihood phylogeny of *pepM* homologs, with bitscores represented by coloured outlines surrounding each branch. Orthologs (orange circles) were identified and selected for further analyses based on their bitscore and relationship to query sequences (yellow stars) from the respective HMM profiles. White circles indicate ultrafast bootstrap support values => 95.

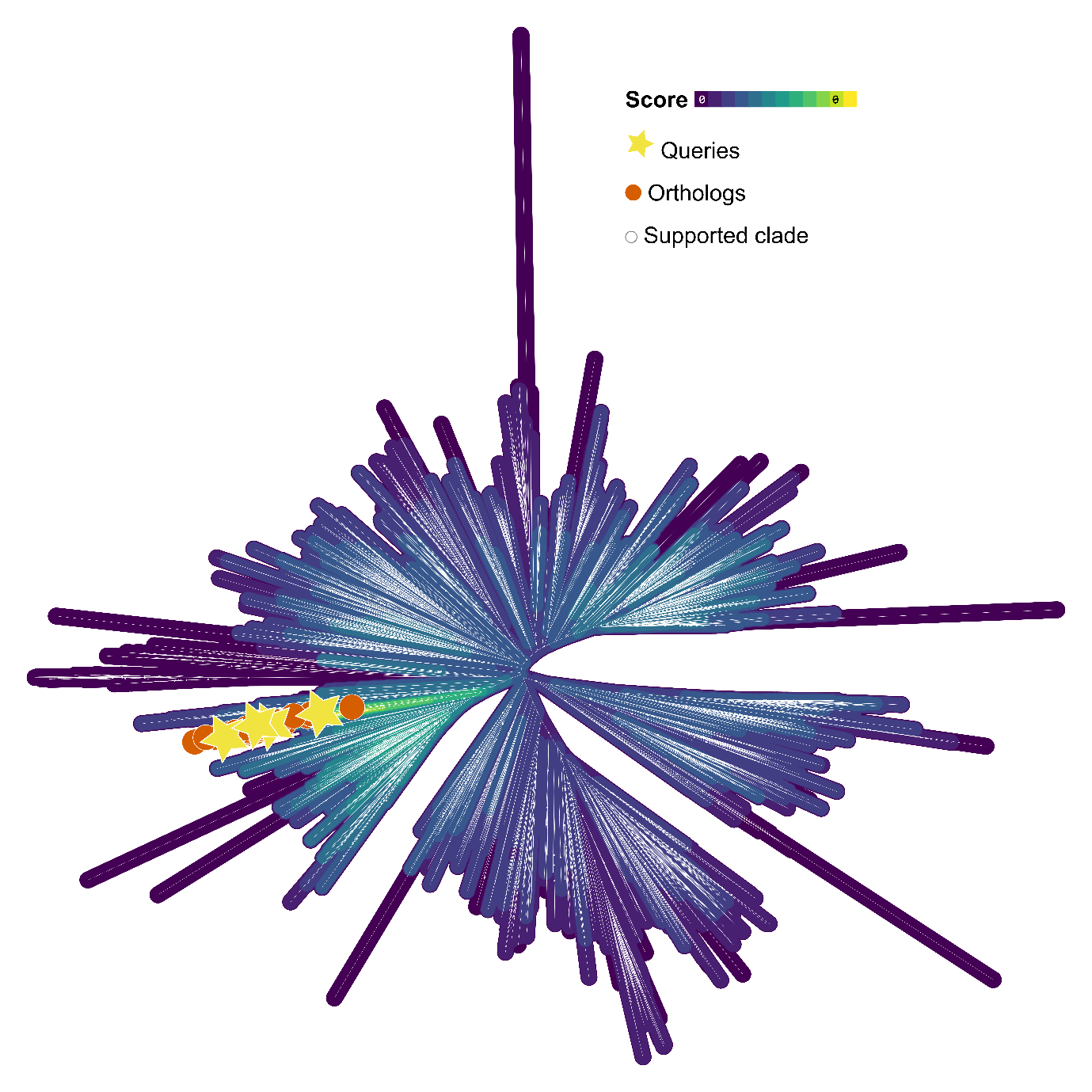

**Figure 2K: Identification of *ptxD* orthologs.** Maximum-likelihood phylogeny of *ptxD* homologs, with bitscores represented by coloured outlines surrounding each branch. Orthologs (orange circles) were identified and selected for further analyses based on their bitscore and relationship to query sequences (yellow stars) from the respective HMM profiles. White circles indicate ultrafast bootstrap support values => 95.

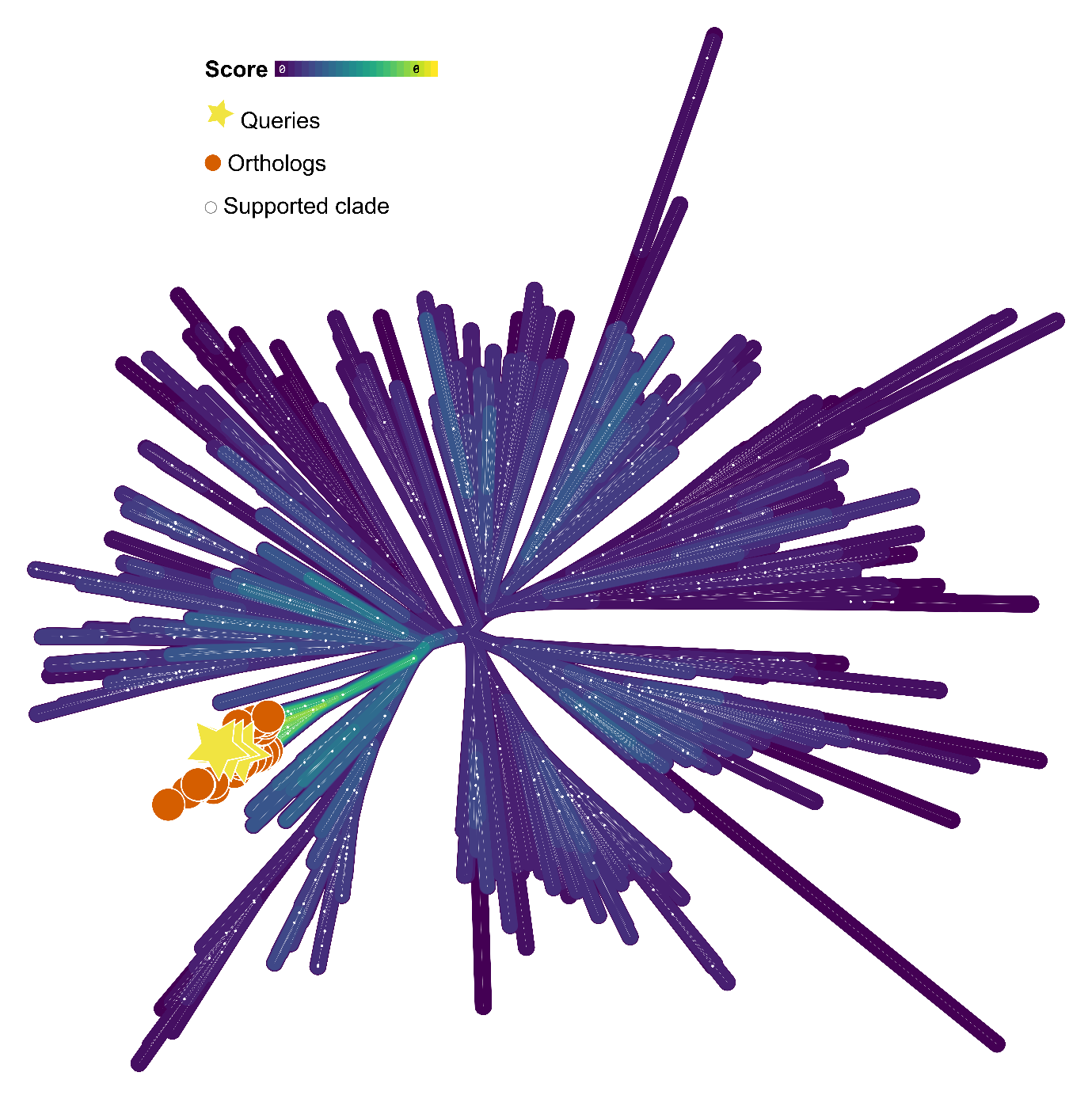

**Figure 2L: Identification of *ptxB* orthologs.** Maximum-likelihood phylogeny of *ptxB* homologs, with bitscores represented by coloured outlines surrounding each branch. Orthologs (orange circles) were identified and selected for further analyses based on their bitscore and relationship to query sequences (yellow stars) from the respective HMM profiles. White circles indicate ultrafast bootstrap support values => 95.

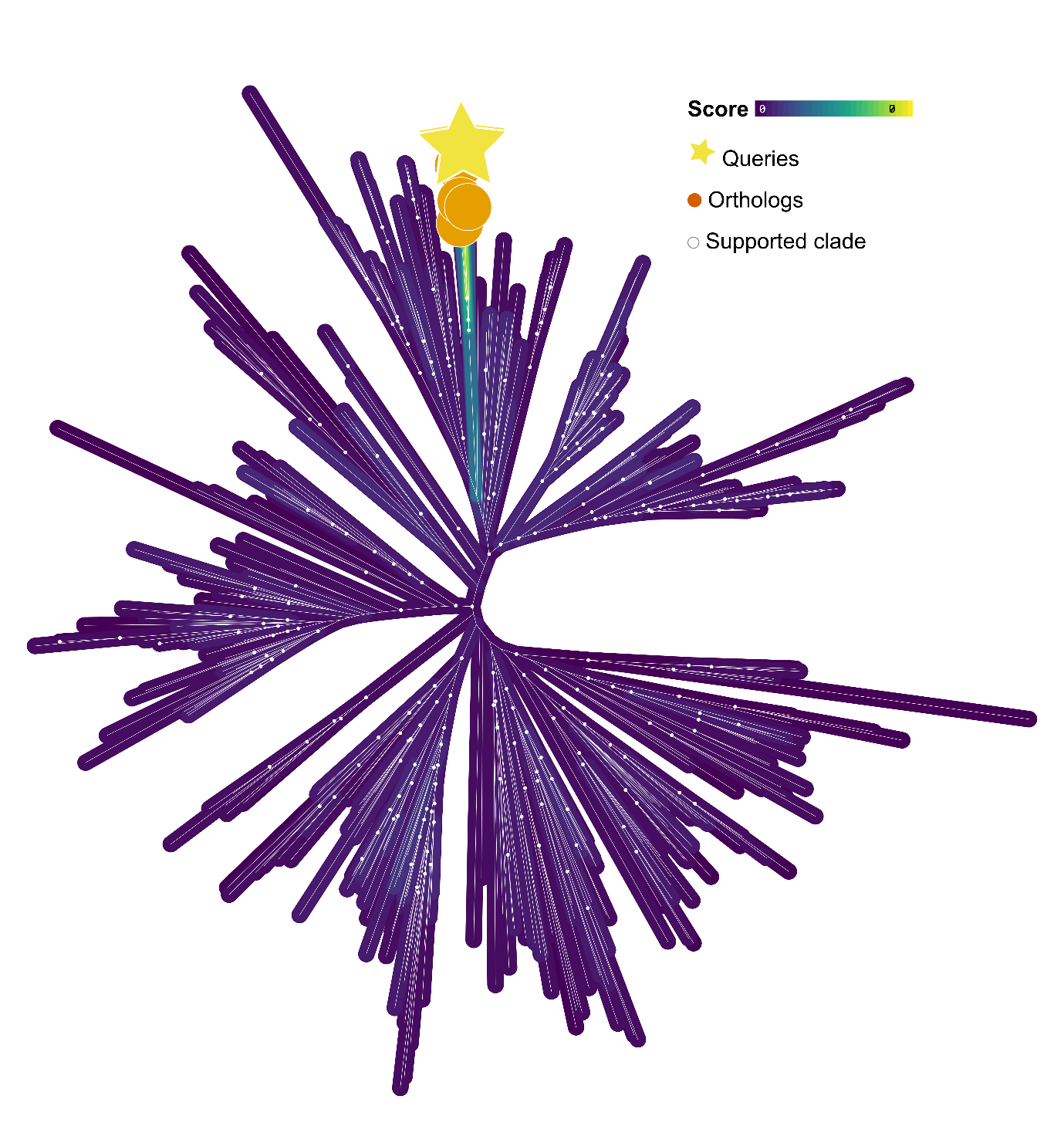

**Figure 2M: Identification of *htxB* orthologs.** Maximum-likelihood phylogeny of *htxB* homologs, with bitscores represented by coloured outlines surrounding each branch. Orthologs (orange circles) were identified and selected for further analyses based on their bitscore and relationship to query sequences (yellow stars) from the respective HMM profiles. White circles indicate ultrafast bootstrap support values => 95.

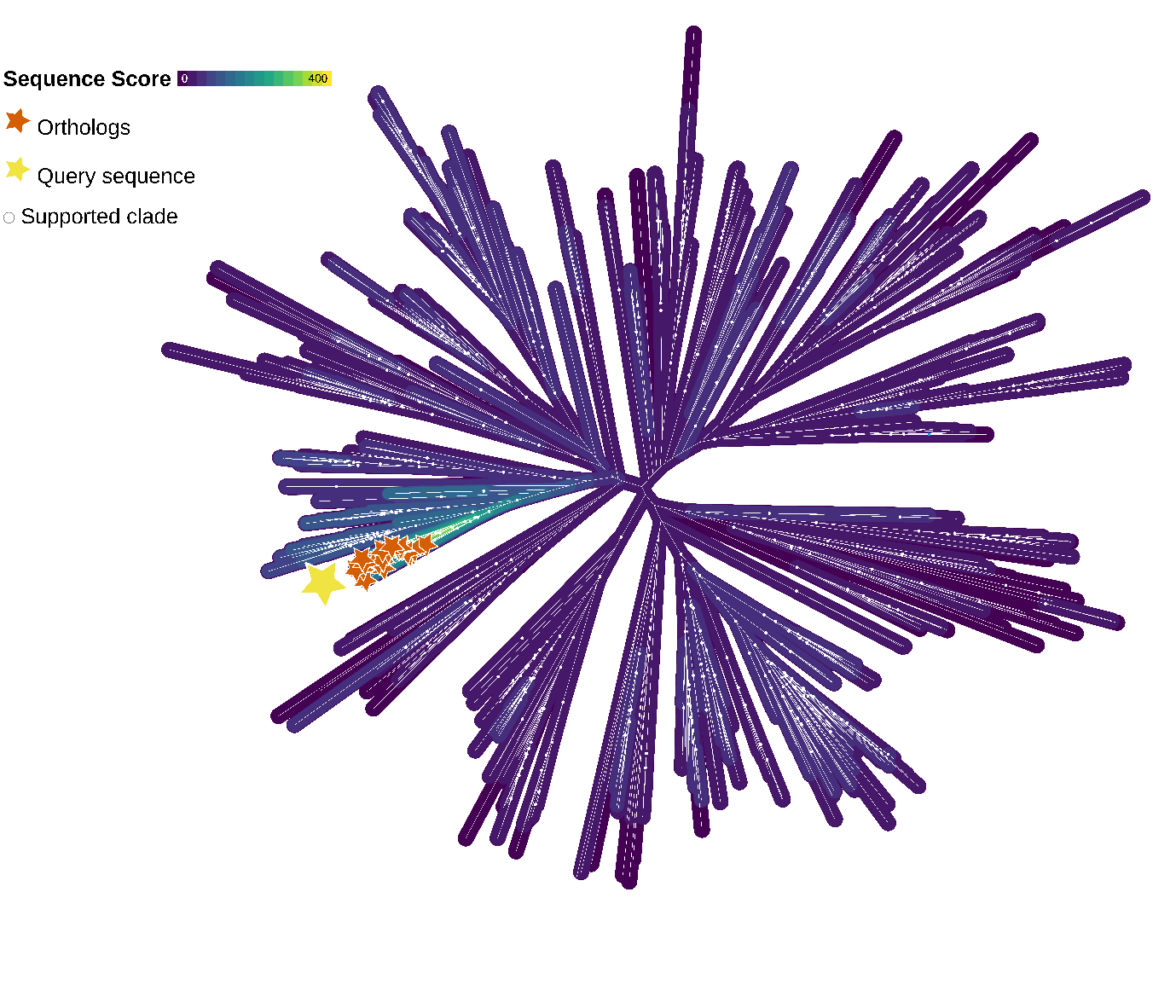

**Figure 2N: Identification of *htxA* orthologs.** Maximum-likelihood phylogeny of *htxA* homologs, with bitscores represented by coloured outlines surrounding each branch. Orthologs (orange stars) were identified and selected for further analyses based on their bitscore and relationship to query sequences (yellow stars) from the respective HMM profiles. White circles indicate ultrafast bootstrap support values => 95.
