## Extended Data for "Timing the Evolution of Phosphorus-Cycling Enzymes Through Geological Time"

**Extended Data Table 1: Phosphorus-cycling genes analysed in this study.** Number of homologs is presented alongside the number of genomes which encode each gene and the number of speciations (spe), horizontal gene transfers (hgt), duplications (dup) and losses (los) predicted by ecceTERA using CIR, LN, and UGAM clock models.

| Phosphorus-Cycling Gene | Homologs | Genomes |  | CIR | | | |  | LN | | | |  | UGAM | | | |
| --- | --- | --- | --- | --- | --- | --- | --- | --- | --- | --- | --- | --- | --- | --- | --- | --- | --- |
|  |  |  |  | spe | hgt | dup | los |  | spe | hgt | dup | los |  | spe | hgt | dup | los |
| *phnJ* | 45 | 45 |  | 28 | 32 | 0 | 18 |  | 31 | 31 | 0 | 19 |  | 27 | 38 | 1 | 21 |
| *phnM* | 69 | 53 |  | 29 | 64 | 4 | 27 |  | 29 | 64 | 3 | 27 |  | 39 | 63 | 5 | 37 |
| *phnZ* | 42 | 40 |  | 39 | 29 | 2 | 26 |  | 38 | 35 | 2 | 29 |  | 38 | 35 | 2 | 28 |
| *phnX* | 29 | 28 |  | 9 | 33 | 1 | 10 |  | 9 | 33 | 1 | 10 |  | 9 | 32 | 2 | 12 |
| *phnA* | 12 | 12 |  | 0 | 11 | 0 | 0 |  | 0 | 11 | 0 | 0 |  | 0 | 11 | 0 | 0 |
| *phnW* | 28 | 25 |  | 8 | 28 | 2 | 8 |  | 8 | 28 | 2 | 8 |  | 8 | 28 | 2 | 8 |
| *ppd* | 36 | 33 |  | 11 | 33 | 3 | 11 |  | 11 | 33 | 3 | 11 |  | 11 | 34 | 4 | 13 |
| *pepM* | 42 | 41 |  | 14 | 47 | 0 | 18 |  | 14 | 47 | 1 | 18 |  | 14 | 42 | 0 | 15 |
| *ptxD* | 50 | 49 |  | 14 | 50 | 3 | 12 |  | 22 | 52 | 2 | 22 |  | 22 | 52 | 2 | 23 |
| *ptxB* | 41 | 38 |  | 22 | 41 | 0 | 19 |  | 18 | 39 | 1 | 17 |  | 18 | 39 | 1 | 17 |
| *htxB* | 5 | 4 |  | 2 | 3 | 1 | 2 |  | 2 | 3 | 1 | 2 |  | 2 | 3 | 1 | 2 |
| *htxA* | 20 | 17 |  | 10 | 19 | 0 | 11 |  | 3 | 18 | 3 | 5 |  | 3 | 18 | 2 | 5 |
| *pitH* | 0 | 0 |  | n/a | n/a | n/a | n/a |  | n/a | n/a | n/a | n/a |  | n/a | n/a | n/a | n/a |
| *pitA* | 1 | 1 |  | n/a | n/a | n/a | n/a |  | n/a | n/a | n/a | n/a |  | n/a | n/a | n/a | n/a |
| *pstS* | 469 | 174 |  | 93 | 210 | 269 | 84 |  | 91 | 212 | 268 | 83 |  | 87 | 212 | 273 | 82 |
| *pnas* | 345 | 257 |  | 171 | 325 | 33 | 163 |  | 176 | 329 | 34 | 176 |  | 176 | 347 | 30 | 186 |

| **Extended Data Table 2: Calibration points used in molecular clock analyses.** We stress that these are conservative estimates, as the origin of a metabolism or group of organisms may predate its first widely-accepted expression in the rock record. | | | | | | |
| --- | --- | --- | --- | --- | --- | --- |
| Calibration | Minimum Age | Citation(s) | Maximum Age | Citation (s) | Phylogenetic Placement | Citation(s) |
| Methanogenesis | 2.7 Ga | ^88^ | n/a | n/a | MRCA of TACK and Euryarchaeota | ^31,89,90^ |
| Red algae | 1.05 Ga | ^91^ | n/a | n/a | MRCA of red algae and red algal chloroplasts | ^78^ |
| Oxygenic photosynthesis | 2.32 Ga | ^92^ | 2.7 Ga | ^93^ | MRCA of photosynthetic cyanobacteria | Cyanobacteria were the first producers of biogenic oxygen |
| Eukaryotes | 1.7 Ga | ^94^ | n/a | n/a | MRCA of Eukaryotes and Archaea | n/a |
| Akinetes | 1.6 Ga | ^95^ | n/a | n/a | First radiation of heterocyst-forming cyanobacteria | ^77^ |
| Prymnesiophyte endosymbionts | 91 Ma | ^96^ | n/a | n/a | MRCA of UCYNA | ^96^ |
| *Hemiaulus* endosymbionts | 110 Ma | ^97^ | n/a | n/a | MRCA of *Richelia intracellularis HH01* and *HM01* | ^98^ |
| MRCA: Most recent common ancestor | | | | | | |

| **Extended Data Table 3: Source of HMM profiles used to identify phosphonate cycling genes**. Each was downloaded from an equivalog HMM used in the NCBI’s prokaryotic genome annotation pipeline. | |
| --- | --- |
| Protein | Equivalog HMM |
| PhnZ | TIGR03276.1 |
| PhnX | TIGR01422.2 |
| PhnA | TIGR02335.1 |
| PhnM | TIGR02318.1 |
| PhnJ | PF06007 |
| PhnW | TIGR02326.1 |
| PepM | TIGR02320.1 |
| Ppd | TIGR03405.1 |
| PstS | TIGR00975.1 & NF008171.0 |
| PNaS | NF037997.1 |
| PitH | NBR010556 |
| PitA | NF03774.1 |

| **Extended Data Table 4: Proteins used to build HMM profiles for PtxD, PtxB, HtxB and HtxA** | | | |
| --- | --- | --- | --- |
| Protein | NCBI ID | Citation | Organism |
| PtxD | YP_001091477.1 | ^18^ | *Prochlorococcus marinus str.* MIT 9301 |
|  | ADB92513.1 | ^24^ | *Desulfotignum phospitoxidans* |
|  | K18916* | ^23^ | *Pseudomonas stutzeri* WM88 |
|  | AAT12779.1 | ^99^ | *Alcaligenes faecalis* WM2072 |
| PtxB | AAC71707.1 | ^18,100,101^ | *Pseudomonas stutzeri* WM88 |
|  | YP_001091475.1 | ^18,101,102^ | *Prochlorococcus marinus str.* MIT 9301 |
|  | ABG49835 | ^101^ | *Trichodesmium erythreaum* IMS101 |
| HtxB | AAC71712.1 | ^101,103^ | *Pseudomonas stutzeri* WM88 |
|  | AAT12776.1 | ^99^ | *Alcaligenes faecalis* WM2072 |
| HtxA | AAC71711.1 | ^39,99,100^ | *Alcaligenes faecalis* WM2072 |

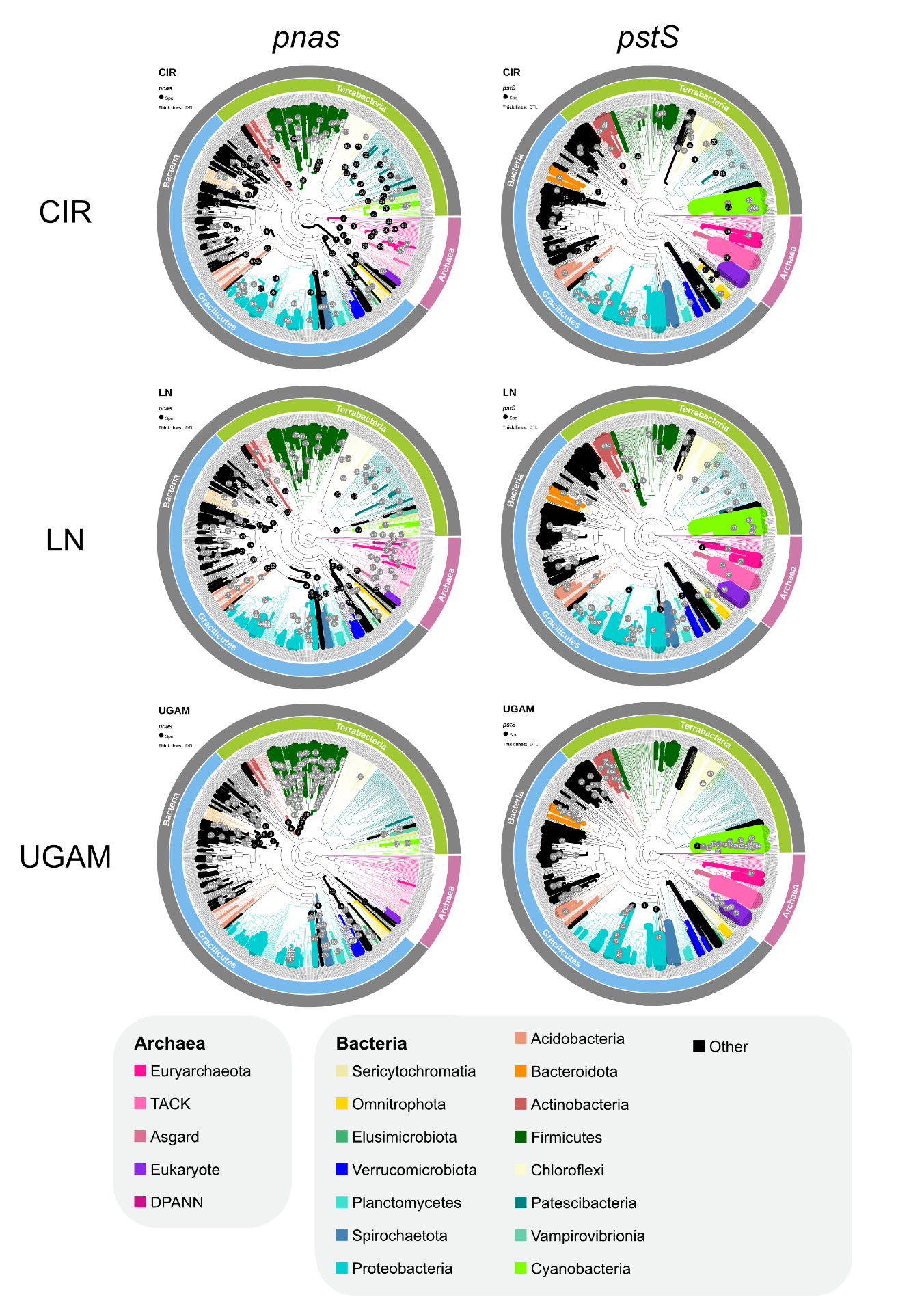

**Extended Data Figure 1: Phylogenetic trees representing where speciations (numbered circles), duplications, transfers and losses (thick lineages) of *pstS* (left) and *pnas* (right) occurred in the tree of life based on three different molecular clock models**. Speciations are numbered in chronological order which Archaean events highlighted in black circles, and others in grey circles. The thickness of each branch corresponds to the number of duplications, transfers and losses that occurred on the branch, whereas branch colours represent which phyla the lineage belongs to.

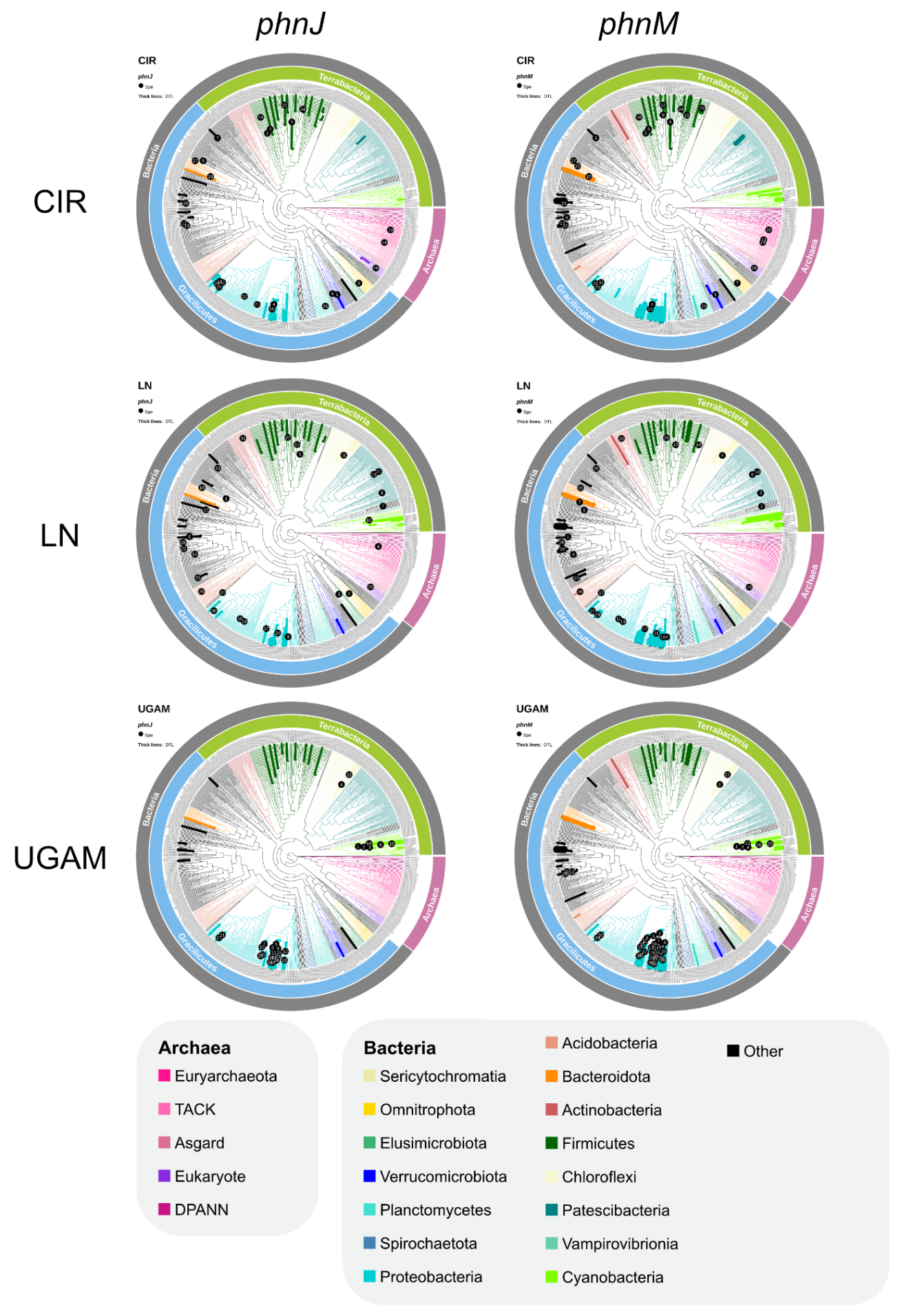

**Extended Data Figure 2: Phylogenetic trees representing where speciations (black numbered circles), duplications, transfers and losses (thick lineages) of *phnJ* (left) and *phnM* (right) occurred in the tree of life based on three different molecular clock models**. Speciations are numbered in chronological order. The thickness of each branch corresponds to the number of duplications, transfers and losses that occurred on the branch, whereas branch colours represent which phyla the lineage belongs to.

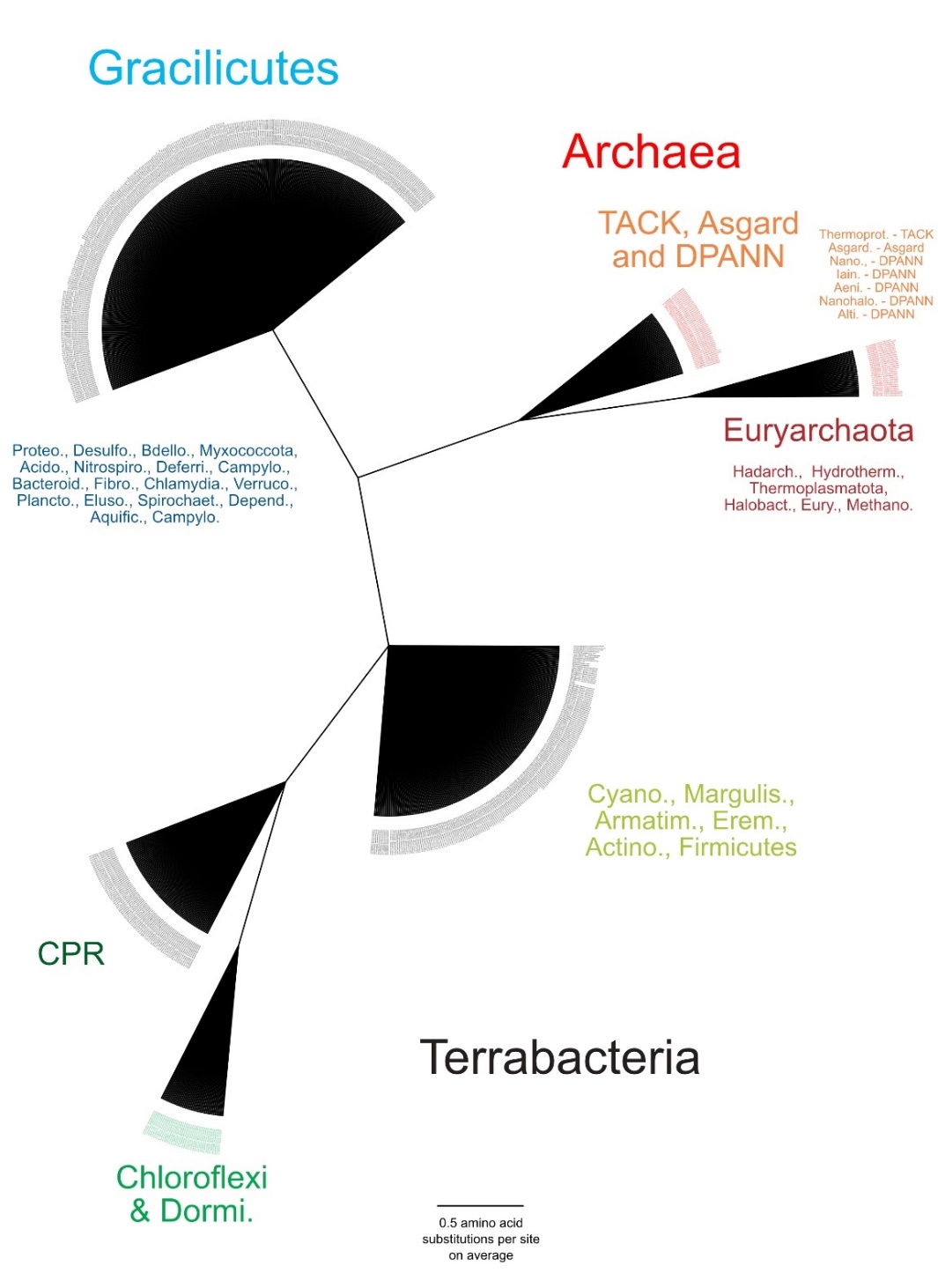

**Extended Data Figure 3: Constraints applied to the tree of life and molecular clock.** Coloured text next to wedges describes which phyla they contain, including Archaea (red and orange), Bacteria of the Gracilicutes clade (blue) and bacteria of the Terrabacteria clade (green). The constraint ensures that Euryarchaeota are more closely-related to each other than they are to other Archaea to match the findings of previous archaeal phylogenies ^31,89^. Similarly, Gracilicutes are constrained to be more closely related to each other than to Terrabacteria or Archaea and vice versa based on the findings of ^29,30,33-36,104,105^. Although previous research (e.g. ^29,30,36,84,106^) produces conflicting evidence on the monophyly of Terrabacteria, we chose to constrain them here because tree certainty metrics have found that artefacts in the form of long-branch attraction towards Archaea and uneven taxon sampling can split the Terrabacterial phylum into separate parts ^33^. CPR are assumed to be closely-related to Chloroflexi and Dormibacteriota within the Terrabacteria despite some conflicting evidence (e.g. ^29,36,84^) because their placement outside of Terrabacteria has also been found to be a result of uneven taxon sampling ^30,33-35^.

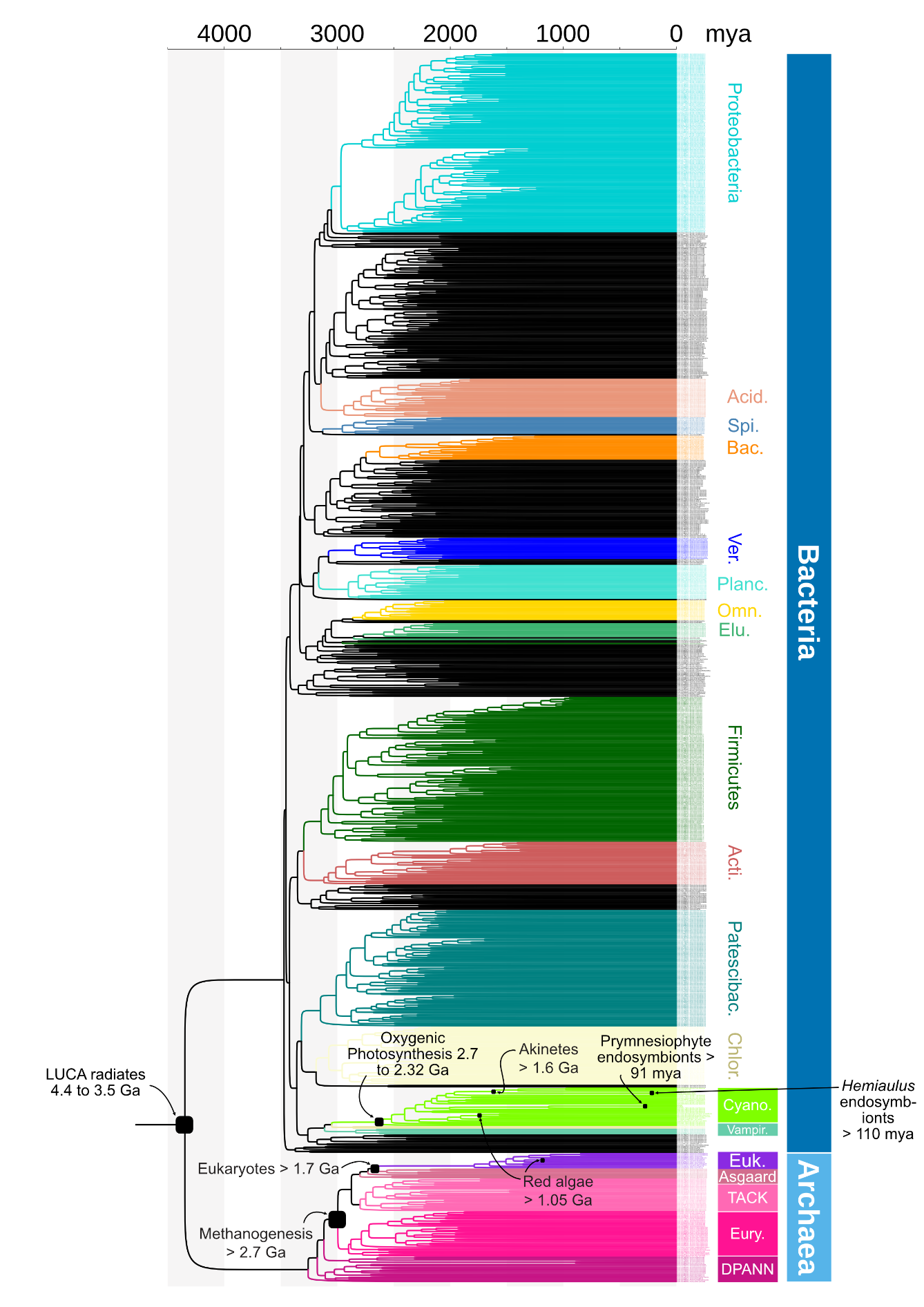

**Extended Data Figure 4: Calibrations for molecular clock analyses.** The position of each calibration point are annotated in black text over a time-calibrated tree made with the CIR clock model. Coloured branches indicate different bacterial and archaeal phyla labelled in text of the same colour. Acid. Acidobacteriota, Spi. Spirochaetota, Bac., Bacteroidota, Ver., Verrucomicrobiota, Planc. Planctomycetota, Omn. Omnitrophota, Elu. Elusimicrobiota, Acti. Actinobacteriota, Patescibac. Patescibacteria, Chlor. Chloroflexota, Cyano. Cyanobacteria, Vampiro. Vampirovibrionia, Euk. Eukaryota, Eury. Euryarchaeota.

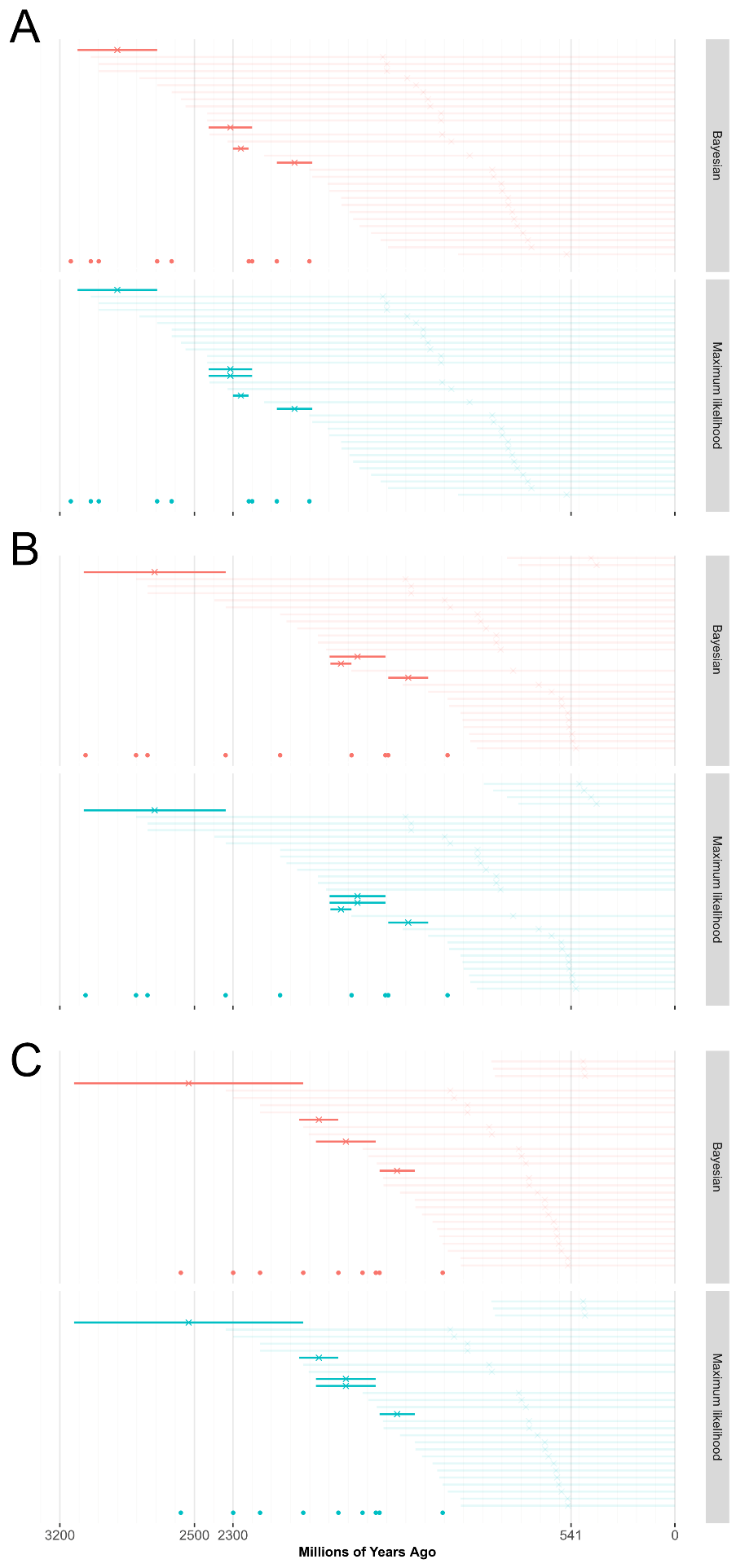

**Extended Data Figure 5: Differences between estimating the origin of *phnX* using cir (A), ln (B) and ugam (c) with Bayesian and Maximum Likelihood methodology .** Bayesian results (orange) were collected using MrBayes to reconstruct the *phnX* phylogeny, whereas maximum likelihood results (cyan) were collected using IQ-TREE to reconstruct the *phnX* phylogeny. Horizontal lines represent the lengths of internal (dark colours) and terminal (pale colours) branches on which gene duplications, transfers and losses are predicted to have occurred.
